# YY1 mutations disrupt corticogenesis through a cell-type specific rewiring of cell-autonomous and non-cell-autonomous transcriptional programs

**DOI:** 10.1101/2024.02.16.580337

**Authors:** Marlene F. Pereira, Veronica Finazzi, Ludovico Rizzuti, Davide Aprile, Vittorio Aiello, Luca Mollica, Matteo Riva, Chiara Soriani, Francesco Dossena, Reinald Shyti, Davide Castaldi, Erika Tenderini, Maria Teresa Carminho-Rodrigues, Julien F. Bally, Bert B.A. de Vries, Michele Gabriele, Alessandro Vitriolo, Giuseppe Testa

**Affiliations:** Department of Experimental Oncology, European Institute of Oncology IRCCS, Via Adamello 16, 20139, Milan, Italy; Department of Oncology and Hemato-Oncology, University of Milan, Via Santa Sofia 9, 20122, Milan, Italy; Human Technopole, Viale Rita Levi-Montalcini 1, 20157, Milan, Italy; Department of Medical Biotechnology and Translational Medicine, University of Milan, Italy; Telethon Institute of Genetics and Medicine (TIGEM), Pozzuoli, Italy; Department of Genetics, University of Geneva & University Hospitals of Geneva, Geneva, Switzerland; Service of Neurology, Department of Clinical Neurosciences, Lausanne University Hospital & University of Lausanne, Lausanne, Switzerland; Radboud University UMC, Nijmegen, The Netherlands; Department of Biological Engineering, Massachusetts Institute of Technology; Cambridge, MA 02139, USA; The Broad Institute of MIT and Harvard; Cambridge, MA 02139, USA; Koch Institute for Integrative Cancer Research; Cambridge, MA, 02139, USA

**Author notes:** Authors contributed equally (Marlene F. Pereira, Veronica Finazzi). Senior authors (Michele Gabriele, Alessandro Vitriolo, Giuseppe Testa).

## Abstract

Germline mutations of YY1 cause Gabriele-de Vries syndrome (GADEVS), a neurodevelopmental disorder featuring intellectual disability and a wide range of systemic manifestations. To dissect the cellular and molecular mechanisms underlying GADEVS, we combined large-scale imaging, single-cell multiomics and gene regulatory network reconstruction in 2D and 3D patient-derived physiopathologically relevant cell lineages. YY1 haploinsufficiency causes a pervasive alteration of cell type specific transcriptional networks, disrupting corticogenesis at the level of neural progenitors and terminally differentiated neurons, including cytoarchitectural defects reminiscent of GADEVS clinical features. Transcriptional alterations in neurons propagated to neighboring astrocytes through a major non-cell autonomous pro-inflammatory effect that grounds the rationale for modulatory interventions. Together, neurodevelopmental trajectories, synaptic formation and neuronal-astrocyte cross talk emerged as salient domains of YY1 dosage-dependent vulnerability. Mechanistically, cell-type resolved reconstruction of gene regulatory networks uncovered the regulatory interplay between YY1, NEUROG2 and ETV5 and its aberrant rewiring in GADEVS. Our findings underscore the reach of advanced in vitro models in capturing developmental antecedents of clinical features and exposing their underlying mechanisms to guide the search for targeted interventions.

## Introduction

YY1 is a ubiquitous zinc-finger transcription factor (TF) whose haploinsufficiency causes Gabriele-de Vries syndrome (GADEVS, MIM: 600013), a predominantly neurodevelopmental disorder featuring mostly intellectual disability, gliosis, hypo- and dystonia with other additional congenital abnormalities of the eye, heart, kidney, genital and skeletal system in a subset of patients^1–7^. The role of YY1 in neurodevelopment was first shown in a *Yy1*^+/-^ mouse model^8^, in which a significant fraction of embryos showed exencephaly, pseudo-ventricles, and asymmetry of the developing brain. YY1 exerts its function by binding to regulatory sequences and recruiting chromatin regulators such as the INO80 chromatin remodeling complex, the p300/CBP histone acetylase, polycomb group proteins, cohesin and condensin, eventually also regulating pre-mRNA splicing^9^. Moreover, we previously showed that *YY1* haploinsufficiency caused a defective gene expression regulation associated with a loss of the histone post-translational modification associated with enhancer activation (H3K27Ac) at YY1-bound enhancers^1^. Shortly after, YY1 was showed to act as an enhancer-promoter (E-P) looping factor^10,11^. For these characteristics, GADEVS can be identified as enhanceropathy^12,13^. However, acute depletion of *YY1* in mouse embryonic stem cells showed a minimal effect of E-P looping and transcription activation^14^. This recent evidence raised a fundamental question about the ability of YY1 in forming and maintaining E-P loops and gene expression. This led us to hypothesize that YY1 regulates the establishment of new E-P rather than their maintenance, in a highly tissue-specific manner. The inability to fully understand the role of YY1 and the reason why its haploinsufficiency causes GADEVS poses a limitation in improving affected individuals’ care. To address these questions and to identify the pathological mechanisms underlying impairments in learning processes and brain abnormalities observed in GADEVS individuals, we derived induced pluripotent stem cells (iPSCs) from multiple individuals affected by *YY1* haploinsufficiency and studied its impact in pluripotency as well as 2D and 3D neuronal models.

## Results

### *YY1* pathotogenic mutations disrupt DNA binding and cause widespread transcriptional downregulation in iPSCs

GADEVS is caused by haploinsufficiency of *YY1*, which is the results of truncating mutations or missense variants in the zinc-finger motifs^1^. To study the molecular dysregulations caused by *YY1* mutations in disease-relevant cell types, we focused on two truncating and one missense mutation in the Zinc-finger domain (GAD01: p.(Lys179*); GAD02: p.(Asp129Ilefs*127)^1^, and GAD03: p.(Cys303Arg)^15^ (fig 1a). We derived iPSCs from three affected individuals’ along with healthy donors’ lymphoblastoid cell lines or skin fibroblasts. The iPSC lines were extensively characterized by immunofluorescence and qRT-PCR for pluripotency markers (fig S1a-c), were confirmed vector-free after approximately 10-12 passages in culture (fig. S1d), and had the capacity to differentiate into derivatives of all three embryonic germ layers (fig. S2). Next, we assessed YY1 protein levels in iPSCs. As expected from truncating mutations, we were able to detect halved levels of YY1 in GAD01 and GAD02 (47.62 ± 12.18%; 61.60 ± 7.79% *vs*. CTL, respectively). In GAD03 (285.20 ± 72.06% *vs*. CTL), we observed instead increased expression of *YY1* transcript (fig. S1g) and protein levels (fig. 1b). As a structural regulator directly controlling gene expression programs, the functional test of YY1 dosage is the validation of YY1 DNA binding across control and mutant lines. We thus performed chromatin immunoprecipitation followed by sequencing (ChIP-seq) to profile YY1 binding sites distributions across our cohort, finding its distribution enriched at promoters and distal intergenic regions, consistent with YY1 known transcriptional regulation function (fig. S3a). Moreover, our ChIP-seq showed coherent peaks distributions with respect to reference ENCODE tracks from H1 human embryonic stem cells (hESC), while YY1 motifs distribution seems equally represented across genomic regions, with a higher abundance at introns and distal intergenic regions (fig. S3b). Finally, the human YY1 motif (MA0095.2 in JASPAR 2020) was by far the most enriched in iPSC YY1 ChIP-seq (fig. S3c), approximately occupying 28% of YY1 peaks in CTLs (*P*< 1 × 10-3311, 2% of background),49% of YY1 peaks in GADEVS (*P*< 1 × 10-2293, 3% of background), and 39% of lost peaks, indicating that lost peaks were effectively bound by YY1, but also that YY1 haploinsufficiency depletes co-factors bindings while retaining high affinity sites. Despite the increased protein levels in GAD03, YY1 ChIP-seq showed a general decrease in binding sites across replicates (fig. 1c) and a comparable drop of binding signals in the three affected lines (fig 1d, S3d). As YY1 was described to work as dimer^10,16^, this observation corroborates the prediction that in the mutant setting only ¼ of the dimers would be functional and hence with an equivalent impact of truncating and missense mutations such as p.(Cys303Arg) that emerged from our assays as a dominant negative (fig S1f). Indeed, ¼ of the average level observed in GAD03 is similar to those observed in GAD01 and GAD02, where only half of the protein level was observed. To further probe this hypothesis, we took advantage of molecular dynamics simulations of pathogenic missense mutations of *YY1*, including the one harbored by GAD03, and evaluated the structural features of the DNA-binding subunits of YY1 in mutants with respect to the wildtype form, finding a reduction of protein stability and DNA-binding ability across all mutants (fig.1e, table S4). Coherently, among differentially expressed genes (DEGs) we observed a widespread down-regulation (585 out of 764 with abs (FC) > 1.5 and FDR < 0.05) in the mutant iPSC lines (fig.1f).

**Fig. 1.**
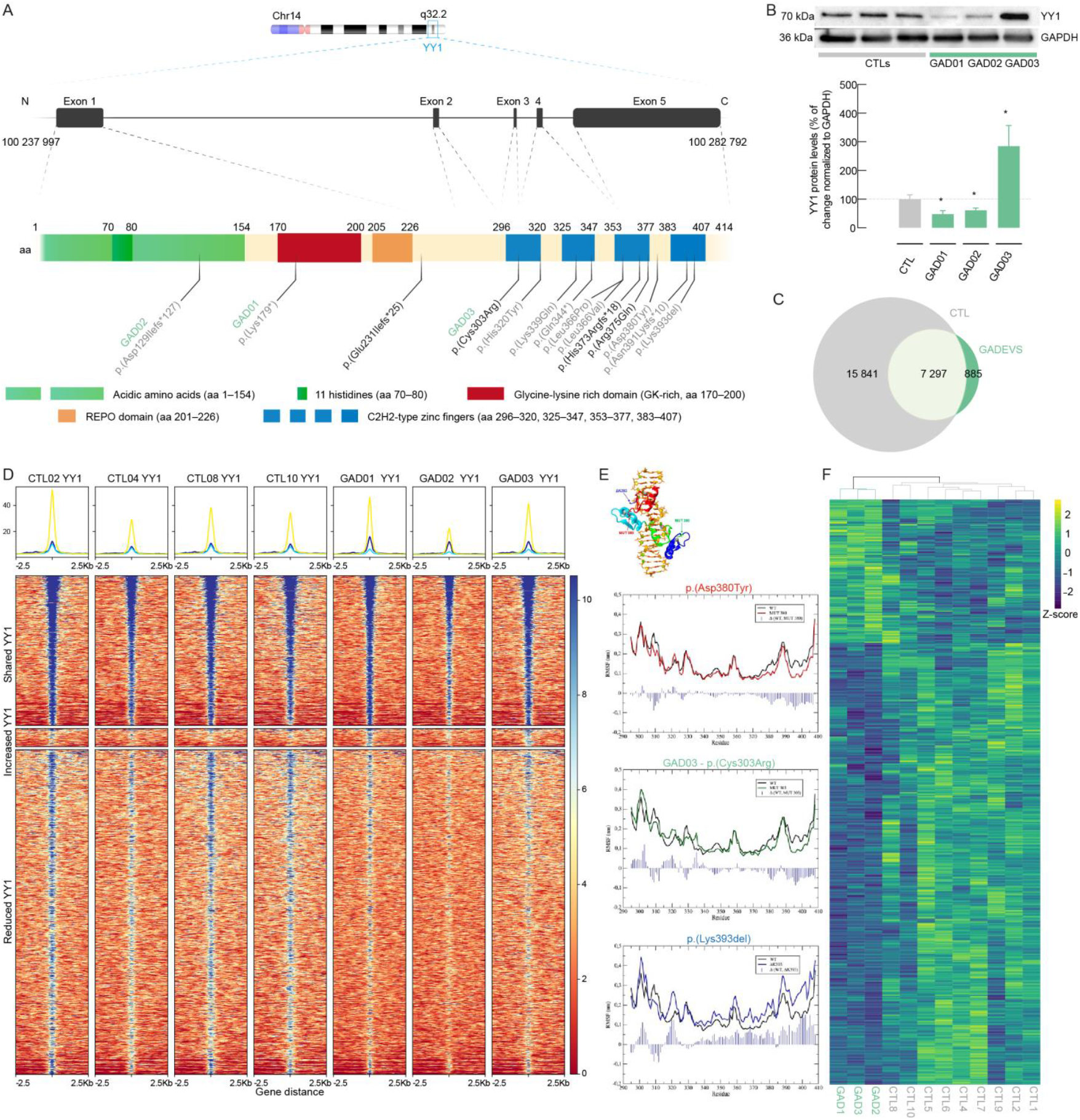
YY1 dosage imbalances are equally detrimental for DNA binding and transcriptional regulation at the pluripotency stage. a) Map of YY1 gene structure and mutation distribution. From the top, YY1 location on chromosome 14 is reported above exon composition and genomic coordinates. Dashed lines report the contribution of each exon to the coded protein. A scheme of the protein is reported with ammino acid residues numbering of each functional domain. Previously described mutations are reported in light grey, newly described mutations are reported in black. On the bottom, a legend describing protein domains is reported. b) YY1 protein levels (normalized to GAPDH) revealed opposite levels of YY1 in GAD03 compared to the remaining GADEVS cohort samples. *P<0.05, *n*=6, unpaired Student’s *t*-test *vs*. CTL. Values are mean ± SEM of *n* independent experiments. *n* refers to the number of protein extracts processed per line. c) Venn diagram reporting the intersection of YY1 peaks in CTL and GADEVS lines. Light grey reports CTL specific peaks (lost), light green refer to shared peaks, dark grey indicates GADEVS specific (gained) peaks. d) Heatmap of library-size normalized YY1 ChIP-seq coverage (RPGC) at YY1 binding sites. Each row represents a 5-kb window centered on peak summits. Reads distribution at Shared, Gained, and Lost peaks are respectively depicted in yellow, blue and cyan. e) Molecular Dynamics of YY1-DNA complex represented in terms of Root Mean Square Fluctuation (RMSF), reported on the y-axis (nm) per residue (x–axis). Flexibility changes are reported around zinc-binding sites. f) Heatmap showing z-scores of log(TMM) read counts for DEGs (764) derived from comparing YY1-mutant vs. control iPSCs (FDR 0.05 and FC 1.5). Yellow and dark blue colors respectively indicate expression levels of up- (179) and down-regulated (585) genes.

### Downregulated YY1 targets enriched for pathologically clinically relevant phenotypes

To understand the impact of the widespread dysregulation observed in affected individuals’ iPSC at the transcriptomic level, we characterized them via enrichment analyses and integration with YY1 ChIP-seq. Gene Ontology (GO) enrichments showed that biological processes relevant for neurological and organogenesis phenotypes associated with GADEVS symptoms were disrupted already at pluripotent stage, as analogously observed in the case of other neurodevelopmental disorders^17,18^ (fig. 2a). Then, we targeted genes directly regulated by YY1 and observed that roughly 25% of downregulated genes were effectively bound by YY1 at regulatory regions (fig. 2b). Notably, among promoters by YY1 we found crucial markers of corticogenesis and, most importantly, genes implicated in a wide range of developmental or neurologic aberrations, such as *SOBP* (associated with mental retardation, retardation with anterior, maxillary protrusion and strabismus (MRAMS) syndrome ^19^), *MLXIPL* (linked with Williams-Beuren syndrome and non-alcoholic fatty liver disease^20^), *KCNN3* (associated with Zimmermann-Laband syndrome^21^), *SEZ6* (occurrence of febrile seizures and dystonia^22,23^), *MAP2* (associated with central neurocytoma within ventricles^24,25^ and neurodegenerative olivopontocerebellar atrophy^26^), *SLC1A1* (implicated in Schizophrenia 18^27^), *OTX2* (syndromic microphthalmia^28–30^), *EEF1A2* (with mutations leading to developmental and epileptic encephalopathy 33, and autosomal dominant intellectual developmental disorder ^31,32^), among many others (fig. 2c). Indeed, Human Phenotype Ontology (HPO) over-representation analysis of YY1 bound down-regulated genes corroborates the observations provided by GO enrichments by highlighting significant enrichments for pathologically relevant phenotypes reported in multiple affected individuals. Among the significant HPOs, we found delayed speech and language development, intellectual disability, seizures, depressed nasal bridge, sensorineural hearing impairment, short stature, visual impairment, and cryptorchidism^5,33^ (fig. 2d).

**Fig. 2.**
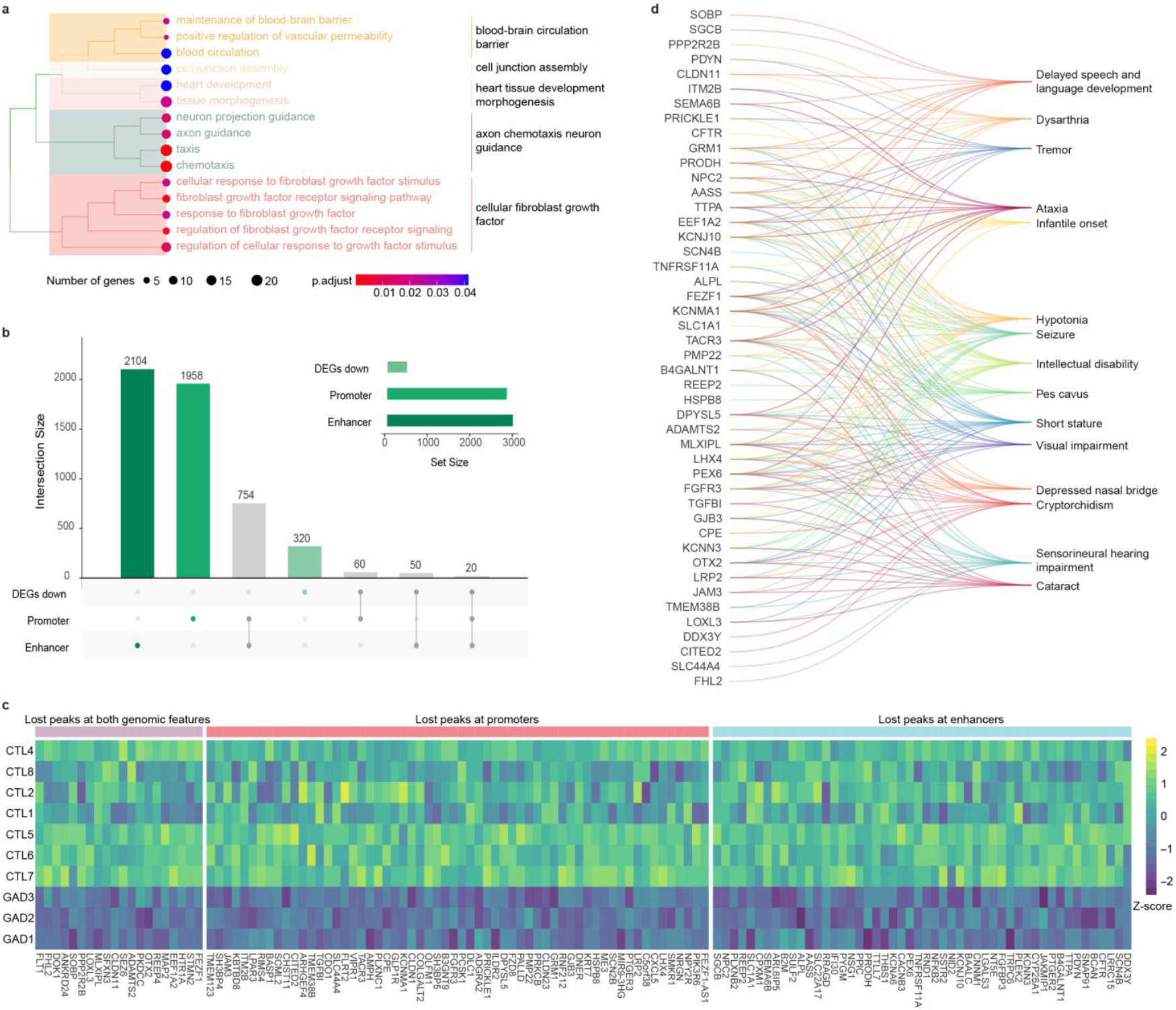
YY1-controlled downregulated genes in iPSC point to pathologically relevant GO and HPO categories. a) Hierarchical clustering representation of GO (biological processes) enrichments. Hierarchical clustering by Jaccard index similarity index is reported on the right. Number of genes defines nodes size and nodes color is defined by p-adjusted of each category enrichment. b) Upset reporting the number of genes whose promoter or enhancers are bound by YY1 in control lines, intersected with down-regulated DEGs. c) Heatmap reporting z-scores of logTMM read counts for down-regulated DEGs, annotated following the intersection reported on panel b. e) Bipartite plot representing gene-HPO associations for differentially expressed genes, and HPO categories significantly enriched by them.

### *YY1* mutations impair the formation of ventricle-like structures at early stages of cortical brain organoid generation

The molecular and cellular origin of GADEVS clinical features remains unexplored. Given the urgency of identifying the molecular alterations caused by *YY1* haploinsufficiency to identify druggable pathways and improve affected individuals’ care, we differentiated iPSCs derived from affected individuals towards clinically relevant cell types. The presence of anomalies in the central and peripheral nervous systems, including intellectual disability, craniofacial and skeletal dysmorphisms, as well as cardiovascular defects, have often been described in GADEVS individuals ^5,33^. Thus, exploring neural crest-derived tissues in *YY1*-mutated samples remains one of the best *in vitro* models to capture craniofacial dysmorphisms. Incidentally, while control lines successfully differentiated into neural crest stem cells (NCSC) (fig. S3e,f), differentiation of iPSCs into NCSCs caused apoptotic blebbing events, similar to what has been already described for the complete loss of *Yy1* in mouse models, that hampers NCSC differentiation^34^, corroborating our observation in iPSC that heterozygous pathogenic mutations of *YY1* induce a loss of function-like phenotype. Since we could not get enough NCSCs for molecular analysis, we resorted to cortical organoids, a complex *in vitro* model representing spatiotemporal neuronal development and capturing the cellular diversity required to understand neurological alterations. To better characterize the pathological relevance of *YY1* mutations in early neuronal differentiation and capture the molecular dysfunction that could explain the mild to severe ID and other neuronal phenotypes such as ventricle atrophy and frontal gliosis observed in GADEVS individuals, iPSCs were differentiated into organoids^35,36^ and cultured for short- (days 3, 7, 12, and 30) and long-term (day 90) assessment (fig. 3a, S3a). First, we sought to assess general morphology properties at early stages of differentiation (fig. S4a) and observed a significant reduction in the cross-section area of spheroid bodies at day 3 (GADEVS *n=*4766, 26438 μm^2^, 427715 *vs*. CTL *n=*8197, 31493 μm^2^, 403003) (fig. S4b-d), mirrored by similar reductions in their perimeter as well (GADEVS *n=*4766, 584.1 μm, 2028 *vs*. CTL *n=*8197, 642.9 μm, 1971) (fig. S4e). These surface imbalances were compensated later, and no further area differences were observed until day 12 (fig. S4f). However, cross-sectional perimeter was increased at day 7 (GADEVS *n*=2722, 984.3 μm, 2723 *vs*. CTL *n*=4578, 952.3 μm, 1868) and 12 (GADEVS *n*=3563, 1405 μm, 5137 *vs*. CTL *n*=8093, 1352 μm, 3881), suggesting that *YY1*-mutated organoids acquired a more irregular shape along differentiation (fig. S4g). To substantiate this hypothesis, we assessed the circularity of control and mutated organoids and indeed, they show a coherent reduction at days 7 (GADEVS *n*=2722, 0.97, 0.29 *vs*. CTL *n*=4578, 0.99 μm, 0.27) and 12 (GADEVS *n*=3563, 0.93, 0.45 *vs*. CTL *n*=8093, 0.99, 0.49) (fig. S4h). Moreover, we calculated the number of organoids in both experimental groups and observed that *YY1*-mutated lines originated less organoids (fig S4i). Regardless of these perimeter and circularity differences, both control and affected cortical organoids showed no clear imbalances in the cellular composition of the progenitor pools (fig. 3b) nor in actively proliferating cells (phospho-histone H3, pHH3-positive cells) within the organoids (fig. S4j). However, following quantification of PAX6-positive ventricle-like structures (VLS), *YY1*-mutated organoids failed to recapitulate the complexity and typical cytoarchitectural organization observed in CTL samples, resulting in significant detection of ‘irregular’ VLS instead. (fig. 3c).

**Fig. 3.**
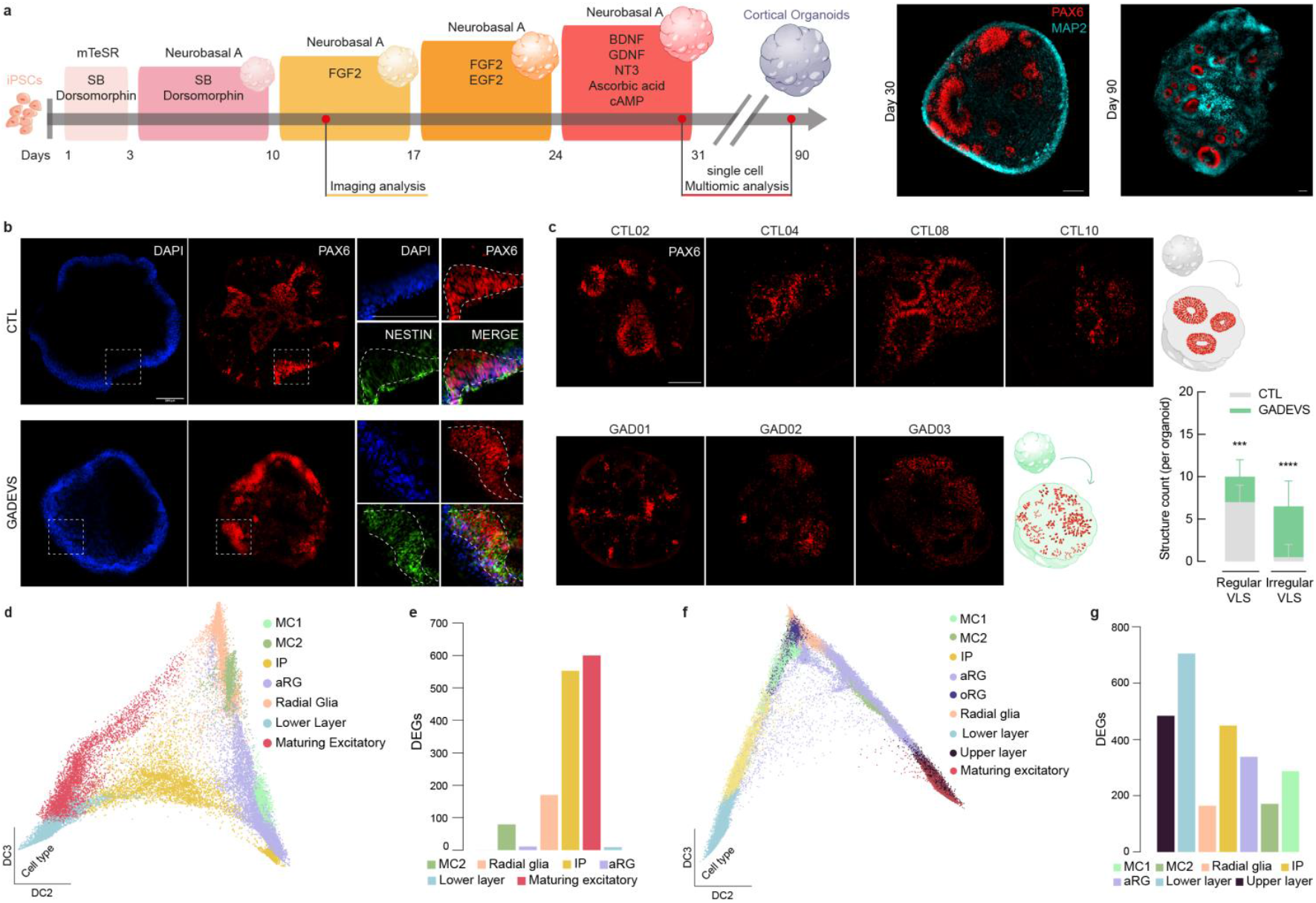
GADEVS cortical brain organoids show impairments in ventricle-like structure (VLS) formation and stage-specific transcriptional disruptions. **a**) Illustration of organoids generation protocol, representing telencephalon small molecule patterning and stage-specific characterization performed. Representative images of immunofluorescence (IF) performed using PAX6, and MAP2 at days 30 and 90, respectively. **b**) Representative pictures of IF and cleared whole cortical organoids with PAX6, and NESTIN for the two different genotypes (CTL and GADEVS). **c**) Representative images of PAX6-positive VLS counted per organoid and classified as regular and irregular depending on the shape and presence of ventricle-like lumen. (whole-organoids acquired in CTL, *n*=32; and GADEVS, *n*=25). Scale bar is 100 μm for all images. **d**) Diffusion map dimensionality reduction reporting cell-type distributions at day 30. **e**) Barplot of differentially expressed genes number per cell-type at day 30. **f**) Diffusion map dimensionality reduction reporting cell-type distributions at day 90. **g**) Barplot of differentially expressed genes number per cell-type at day 90. Scale bar of 100 μm for all images.

### *YY1* haploinsufficiency affects intermediate cellular states in early organoid development and impacts differentiated lineages in later stages

To further characterize the cellular complexity and composition of *YY1*-mutated organoids at day 30 and day 90 regarding gene expression and DNA regulatory regions, we resorted to single cell multiomic profiling. Indeed, single-cell multiome captures the transcriptome and chromatin accessibility within the same cell, allowing the identification of gene-regulatory networks (GRN) along differentiation trajectories^37–39^.

To assign cell-type identities to our datasets, we combined Leiden clustering with the identification of key markers and differentiation trajectories (fig. 3d,f, S5a,b see Methods). To further validate the cell-type annotation, we ingested the two datasets onto a reference single-cell fetal cortex dataset^40^ (fig. S5c,d). Then, to corroborate our cell-type annotation, we intersected the top expressed genes in our clusters with the top expressed in each cell type identified in the reference fetal data (fig. S5e,f). Already at day 30 we identified different types of progenitors, including SOX2^+^ apical radial glia (aRG) like-cells, SLC1A3^+^ radial glia (RG), and EOMES^+^ intermediate progenitors (IP) (fig. 3e). Moreover, we observed two main differentiation paths, mirroring direct (IP-independent) and indirect (IP-dependent) neurogenesis (fig. S5g). We respectively identified i) a cluster of maturating neurons (DCX^+^) derived from a RG cluster, and ii) a cluster of TBR1^+^ cells derived from the IP, on two separated paths (fig. 3d, fig. S5a). Despite detecting irregular VLS at day 12, we did not observe significant differences in cell-type abundances in the RNA modality of our single cell datasets. However, we observed a severe transcriptional dysregulation in RG, IP and maturing excitatory neurons (fig. 3e), which might indicate an impairment in the generation of mature neurons later in development. As expected, at day 90 we identified more cell-types representative of a more mature stage of differentiation, comprising HOPX^+^ outer radial glia (oRG)-like cells, SATB2^+^ upper layer-like neurons, and a minor representation of AQP4^+^ cells, unmasking the presence of a rare population of astrocytes, and at the beginning of gliogenesis (fig. S5b). Of note, at this later stage of development we saw a wide transcriptional dysregulation affecting the majority of identified cell-types (fig. 3g), with neurons derived from both IP and RG as the most disrupted, suggesting a cumulative/epigenetic effect.

### *YY1* haploinsufficiency triggers an extensive rewiring of cell-type specific transcriptional programs

Next, we measured the direct effect of YY1 haploinsufficiency on the underlying GRNs in cortical organoids at day 30. To reconstruct the GRNs in the identified cell-types, we applied Cell Oracle^41^. First, we used GRNs to further confirm the identity of IP by looking at their most central TFs (Fig. S6a). Among the first 30 TFs, we identified migration genes such as *NFIB* and *POU3F2*, and *RORB* in accordance with IP role in originating lower layer neurons. Moreover, we found *EOMES* centrality enriched in IP compared to RG (fig. S6b). Finally, we identified a higher specificity for EOMES targets in the indirect differentiation lineage (Fig. S6c), corroborating that IP-mediated differentiation is better explained by EOMES regulatory activity. Once we validated IP identity, we identified DEGs directly regulated by YY1 in each cell-type. Enriched categories among DEGs directly regulated by YY1 in RG referred to *cell growth* and *positive regulation of developmental process* (fig. 4a, table S3). In IP, direct targets of YY1 enriched *tissue morphogenesis* and related categories such as *neural tube morphogenesis* (fig.4b, table S3). Moreover, direct targets of YY1 significantly enrich DEGs (p = 3.60e-65 for RG and p = 3.789e-95 for IP). Hence, both RG and IP categories enriched by DEGs directly regulated by YY1 point to defects in development, positioning YY1 upstream of the dysregulations observed in both cell-types, and potentially pinpointing the pseudo-ventricle phenotype observed at earliest stages to a core of genes directly controlled by YY1.

**Fig. 4.**
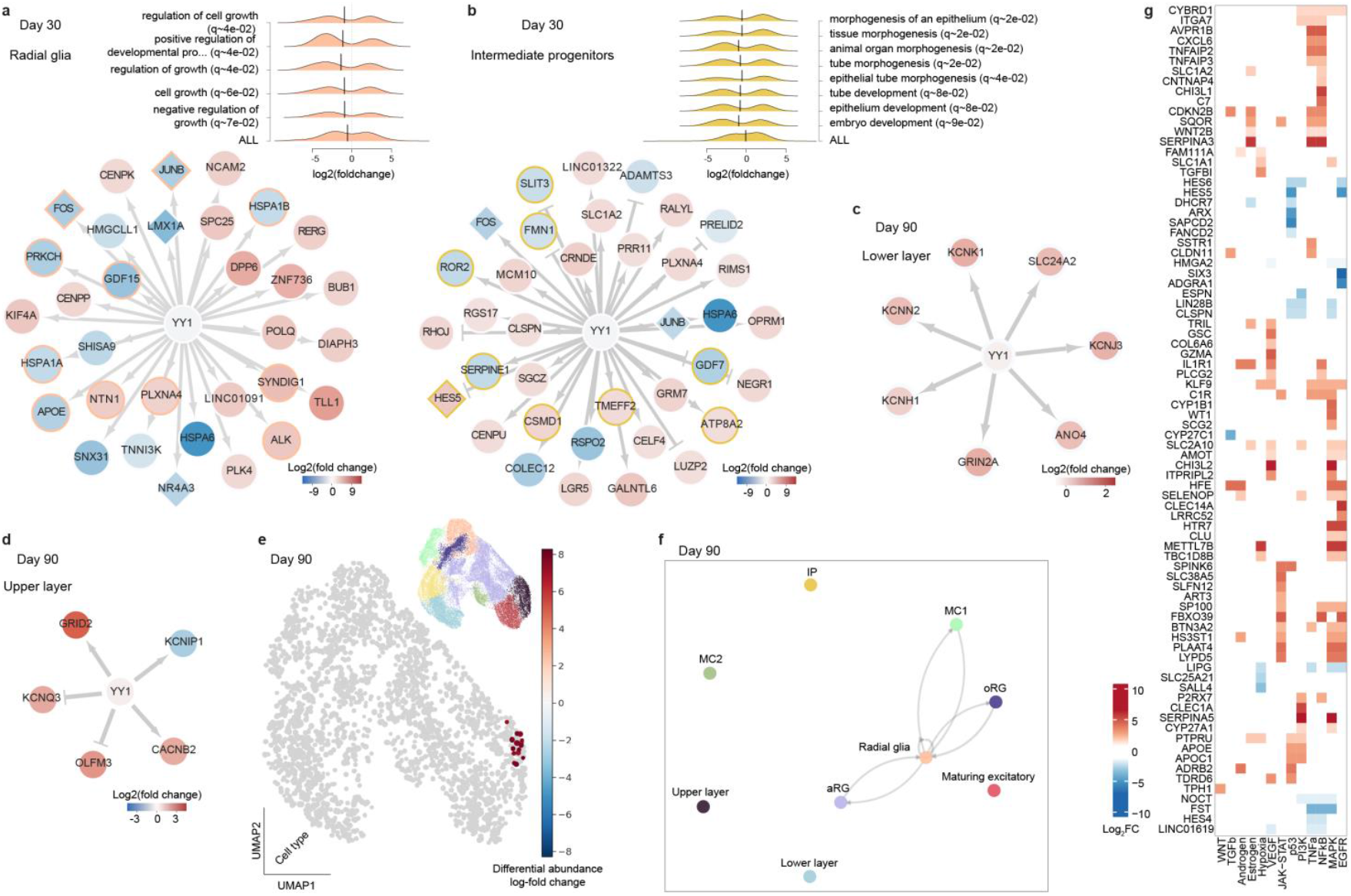
Mechanistic dissection of cell-type specific YY1-dependent dysregulations pinpoint YY1 regulatory activity to cell migration in IP and ion channels in mature neurons, as well as cell-cell communication defects among glia cells in late stages of GADEVS organoids corticogenesis. a) Graph representation of the RG gene-regulatory-network comprising YY1 and its differentially expressed direct targets. Transcription factors are represented as diamond. Nodes are coloured by log2FC. Purple borders represent association to “positive regulation of developmental process” GO category. Arrows represent activation by YY1 and T-shaped edges represent putative repressive activity by YY1. b) Gene regulatory network representation of YY1 direct targets in IP. Arrows represent activation by YY1 and T-shaped edges represent putative repressive activity by YY1. Nodes color represent log2(FC) measured by differential expression on IP pseudobulks, comparing controls with GADEVS lines. Green borders represent association to “morphogenesis” GO categories. c) Gene regulatory network representation of YY1 direct targets enriching cell ion channel category in lower layer-like neurons. Edges and nodes are depicted as in panel b, using log2(FC) from differential expression performed on lower-layer neurons d) Gene regulatory network representation of YY1 direct targets enriching cell ion channel category in upper layer-like neurons. Edges and nodes are depicted as in panel b, using log2(FC) from differential expression performed on upper-layer neurons. e) UMAP of day 90 organoids coloured by differential abundance scores measured with milopy (see methods). A small UMAP coloured by cell-type is reported in the upper right corner for reference. Cell-type colors are the same used in panel f. f) Differential cell-cell interactions calculated by Liana at day 90. Arrows depict interactions between cell-types inferred by expression of receptor-ligand pairs. Only significantly affected interactions are depicted. g) heatmap depicting log2(FC) of genes involved in PROGENy cell-cell signaling pathways, derived from differential expression analysis on pseudobulk of RG ad day 90.

The two mature neuronal subtypes were the two most disrupted cell types at day 90. TBR1^+^ neurons showed disruption in several “*synaptic signalling*” categories, as well as “*long-term synaptic potentiation”*, which are directly connected with neuronal activity and *cholesterol transport*, essential for neuronal metabolism^42,43^ (table S3). Direct targets of YY1 identified as actively regulated in these neurons enrich for DNA binding and categories linked to transcription regulation (table S3). When we focused on differentially expressed YY1 direct targets, we found enrichments for molecular functions related to potassium, calcium and glutamate channel activity, ultimately leading to putative direct impairments on basal synaptic transmission. The same genes enriched cellular components for “*ion channel complex”* and “*synaptic membrane”* (fig. 4c, table S3). SATB2^+^ neurons showed dysregulations in “*synaptic signaling”*, “*neuron development*”, and “*regulation of membrane depolarization”*, which hinted at functional dysregulations (table S3). Notably, “*regulation of cell development”*, “*regulation of synaptic plasticity*” and “*regulation of the nervous system process*” were enriched among YY1 targets expressed in the same neurons (table S3), hinting at a direct link between YY1 activity and the dysregulation globally found in this cell-type. Similarly to what we observed in TBR1^+^ neuronal populations, crossing YY1 targets with DEGs in SATB2^+^ neurons, we found enrichments for cellular compartment categories linked to ion channels, ultimately pinpointing YY1 contribution to their dysregulations in both neuronal lineages through two different gene modules (fig. 4d, table S3).

Since YY1 was not differentially expressed in IP at day 30 and in neurons at day 90, but we found i) cell signaling differentially in IP at day 30, ii) upregulation of YY1 direct targets enriching GO categories related to *gene expression regulation*, and iii) several cell-types that showed broad transcriptional dysregulation at later stages, we speculated that compensatory mechanisms, including non-cell-autonomous effects, might contribute to organoids transcriptional phenotypes. Thus, we investigated cell-type abundances and cell-cell interactions. We observed an increased relative abundance of GADEVS cells in the upper-layer neurons cluster only at day 90 (Fig. 4e). Hence, we quantified^44^ the contribution to gene expression of specific subgroups of receptor-ligand interactions (Fig. S6d) and identified a significant difference in one of the factors (Factor 6) recapitulating cell-cell interactions (fig. S6e). This factor highlights a disruption in the communication centered on RG and linking aRG, oRG, and cycling cells (MC1) clusters (fig. 4f). Finally, we pinpointed the dysregulations enriching the most disrupted cell-cell interaction pathways (fig. 4g). First, we observed a strong enrichment of EGFR and MAPK in Factor 6, and targets of both pathways enriching YY1 targets (p < 0.05) in MC1 and the central RG clusters. However, we did not find a significant enrichment for EGFR or MAPK targets among DEGs of both cell-types, which point to a more subtle but broad dysregulation. Given the non-cell autonomous type of dysregulation observed, and the fact that we found cells expressing AQP4 – an astrocytes-specific water channel - within the central RG cluster (fig. 3e), we speculated that astrocytes-mediated non-cell autonomous functions might also be impaired.

### Abnormal neuronal organization and synaptic impairments in *YY1*-mutated glutamatergic neurons

Given the dysfunctions observed in late stage organoids in lower- and upper-layer neurons, we further probed their mechanistic and functional impact in NGN2-derived glutamatergic neurons (fig. S7a). Given the disruption of non-cell-autonomous interactions in late-stage organoids, and the critical role of astrocytes in neuronal maturation, and response to environmental signals and synapse modulation, *NGN2*-derived neurons were co-cultured with fresh mouse astrocytes using transwell systems in the presence of healthy fresh mouse astrocytes (fig. S7b). This approach enabled us to keep physically separated NGN2-derived neurons from astrocytes while preserving their interaction (fig. 5a). First, bulk RNA-seq in glutamatergic neurons allowed us to quantify allelic abundances of *YY1* (fig. S8b) and observe that *YY1* expression patterns were similar to the ones observed in iPSCs (fig. S8a). Moreover, GAD03 increased expression was coupled to a higher detection of reads from the mutated allele (fig. S8a), further corroborating the dominant negative model emerging from the YY1 loss of function described in iPSC (fig. S1f).

**Fig. 5.**
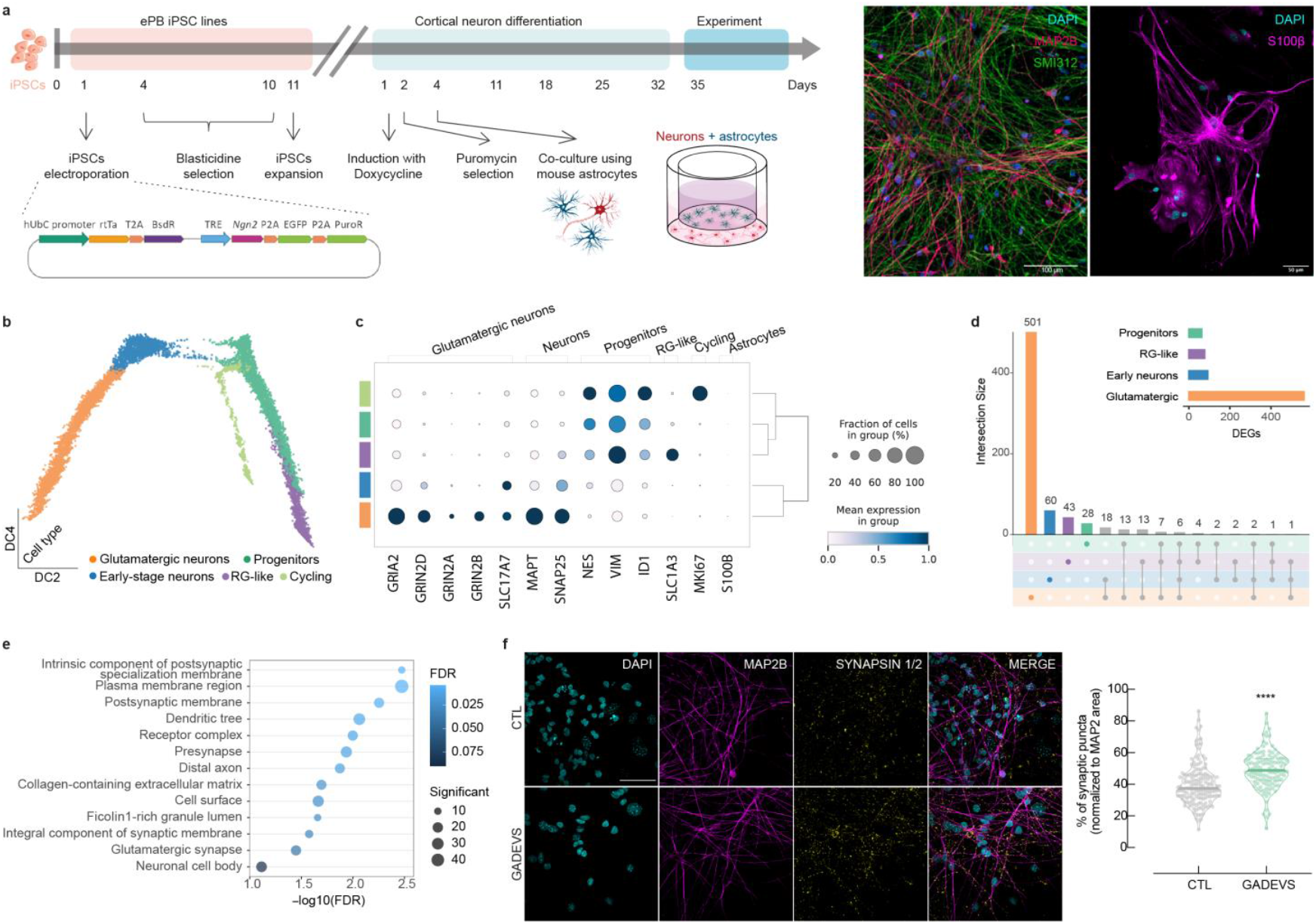
GADEVS iPSC-derived glutamatergic neurons display transcriptional and morphological synaptic abnormalities. **a**) Differentiation protocol to generate glutamatergic neurons starting from human derived iPSC. To induce NGN2 ectopic expression a ePiggyBac plasmid was transfected by electroporation. After selection, iPSC were co-cultured on transwell with fresh mouse astrocytes for 35 days. Immunostainings showed coherent expression of dendritic and pan-axonal markers MAP2B and SMI312 (scale bar 100 μm) and co-culture was achieved by addition of astrocytes (S100β-positive, scale bar 50 μm). b) Diffusion map of single-cell RNA modality from NGN2 iPSC-derived neurons show an heterogenous set of cell-types including early and mature neuronal lineages. A color legend depicting cell-types is provided on the bottom of the panel c) Dotplot showing representative markers of glutamatergic neurons, pan markers of neuronal lineages, and markers of progenitors and glial cells. d) Upset plot reporting the intersection of lists of differentially expressed genes (DEGs). Glutamatergic neurons (orange) show the largest cell-type specific set of DEGs e) Dotplot of GO enrichments (biological process) for genes differentially expressed in the glutamatergic neurons cluster. f) Representative figures of glutamatergic neurons immunostained with MAP2B and synapsins1/2. Scale bar is 50 μm. Synaptic puncta was quantified by counting the number of synapsins 1/2 when co-localized with MAP2B de-noised signal for all fields of view acquired. Graph on the right shows the calculated percentage of synaptic puncta normalized to MAP2B (grouped analysis CTL *n*=156 *vs*. GADEVS *n*=117 fields of view across 4 independent seedings). results are presented as median (min, max, and range) of percentages of synaptic puncta counted in *n* different fields of view independently acquired. \**P*<0.05, \*\*\*\**P*<0.0001 *vs*. CTL using a two-tailed Mann-Whitney test.

To capture the diversity of neuronal fates generated by the comparatively homogeneous *NGN2* neuronal differentiation paradigm ^45^ we applied single cell approaches using gene expression and chromatin accessibility modalities, and we took advantage of the physical separation of the two cell-types to also contextually profile neurons via single cell multiomics and astrocytes via bulk RNA-seq. The single cell transcriptome of neurons grown in co-culture with fresh astrocytes showed the presence of multiple progenitors (fig. S8c, d). Diffusion maps effectively highlighted a population of cycling cells that differentiated into progenitors, which then generated two lineages (fig. 5b), one of *NEUROG2*^+^ *bona fide* neurons, as expected, and surprisingly, a second one of *SLC1A3*^+^ cells (fig.5c, fig. S8c, d), marking a cluster of radial glial-like cells that had not been annotated in previous studies. Additionally, we could identify two different states of neuronal maturation, which we designated as early-stage and glutamatergic neurons (fig. 5c). After performing differential expression analysis (DEA) in cell type-specific clusters, we observed that glutamatergic neurons were the most dysregulated (fig. 5d). Accordingly, GO analysis highlighted significant enrichments for cellular component categories related with glutamatergic pre and postsynaptic compartments (fig. 5e) and to biological processes related to neuronal functions, such as *anterograde trans-synaptic signaling* (fig. S8f), whose disruption was detected also in organoids (table S3). Thus, to account for disparities in synaptic assembly and organization we quantified a group of synapsins, which are important components of mature synapses in the cortex. As anticipated by GO enrichment analysis, *YY1*-mutated neurons exhibited a significant increase of dendritic synaptic puncta (GADEVS *n*=117, 48.83%, 72.70 *vs*. CTL *n*=156, 37.43%, 74.97) (fig. 5f).

### Astrocytes respond to environmental changes caused by *YY1*-deficient neurons and undergo cell activation

Based on the extensive alteration of the synaptic compartments, we reasoned that the consequences of *YY1* haploinsufficiency could extend beyond the affected neurons and adversely affects the physiology of surrounding astrocytes. To measure the non-cell autonomous effect of altered neurons on surrounding cells, we examined the transcriptomic profile of healthy mouse astrocytes in contact with healthy or GADEVS NGN2-derived neurons. We identified 291 DEGs, predominantly upregulated (fig. 6a, 248 and 43 up and downregulated, respectively), which were enriched in GO processes involved in immune regulation and leukocyte reactive responses (fig. 6a). Consistent with how dysfunctional synaptic activity suffices to induce astrogliosis-like responses^46,47^, we observed that all genes enriching these GO categories were found to be exclusively upregulated (genes are depicted in each node in the gene-concept network plot, fig. 6a). To verify the hypothesis that astrocytes are undergoing the activation process marked as astrogliosis, we also performed a network reconstruction using the total significant DEGs (291 genes) to dissect the connectivity of each gene and expose the most critical regulators as means to highlight the central genes. As a result, we identified 193 nodes that formed one complex network in which *Spi1, Myb, Fosb, Foxn4, Neurog2*, and *Tnf* represent master regulators, controlling directly or indirectly most of the nodes (fig. 6b). Of particular interest, *Tnf* (one of the genes with highest degree score in the network) encodes a multifunctional proinflammatory cytokine belonging to the tumor necrosis factor (TNF) superfamily, which has been shown to be one of the main pro-inflammatory cytokines involved in regulation of astrocytes and microglia activation^48,49^. Directly linked with *Tnf*, other TNF ligands and receptors including *Tnfsf9, Tnfrsf11a*, and Tnfrsf25 are among the upregulated DEGs linked with astrogliosis gene signatures. Likewise, macrophage-associated genes (*Fcer1g, Fcgr3, Mpeg1, Lcp2, Rac2, Trem2*, and *Nckap1l*), chemokines, interleukin receptors, channels, and serpin protease inhibitors (*Il1rn, Il7r, Ccl4, Ccl6, Ccl9, Serpinb1*, and *Serpinb9b*), potentiators and markers of astrogliosis (*S100β, Lgals3, Nfkbid, P2ry10b, P2ry12, Nfkbid, Spp1, Aif1, Cd83, Cd1, Pdgfb*, and *Vegfa*), as well as other pivotal genes associated with inflammatory responses (*A2m, Lilrb4, Adam8, Lpl, Slfn2, Plaur, Plau, Cxcl2, Adamtsl4, Hpgds*) are depicted in the DEGs network underscoring a primary composition of astrocyte activation genes (fig. 6b).

**Fig. 6.**
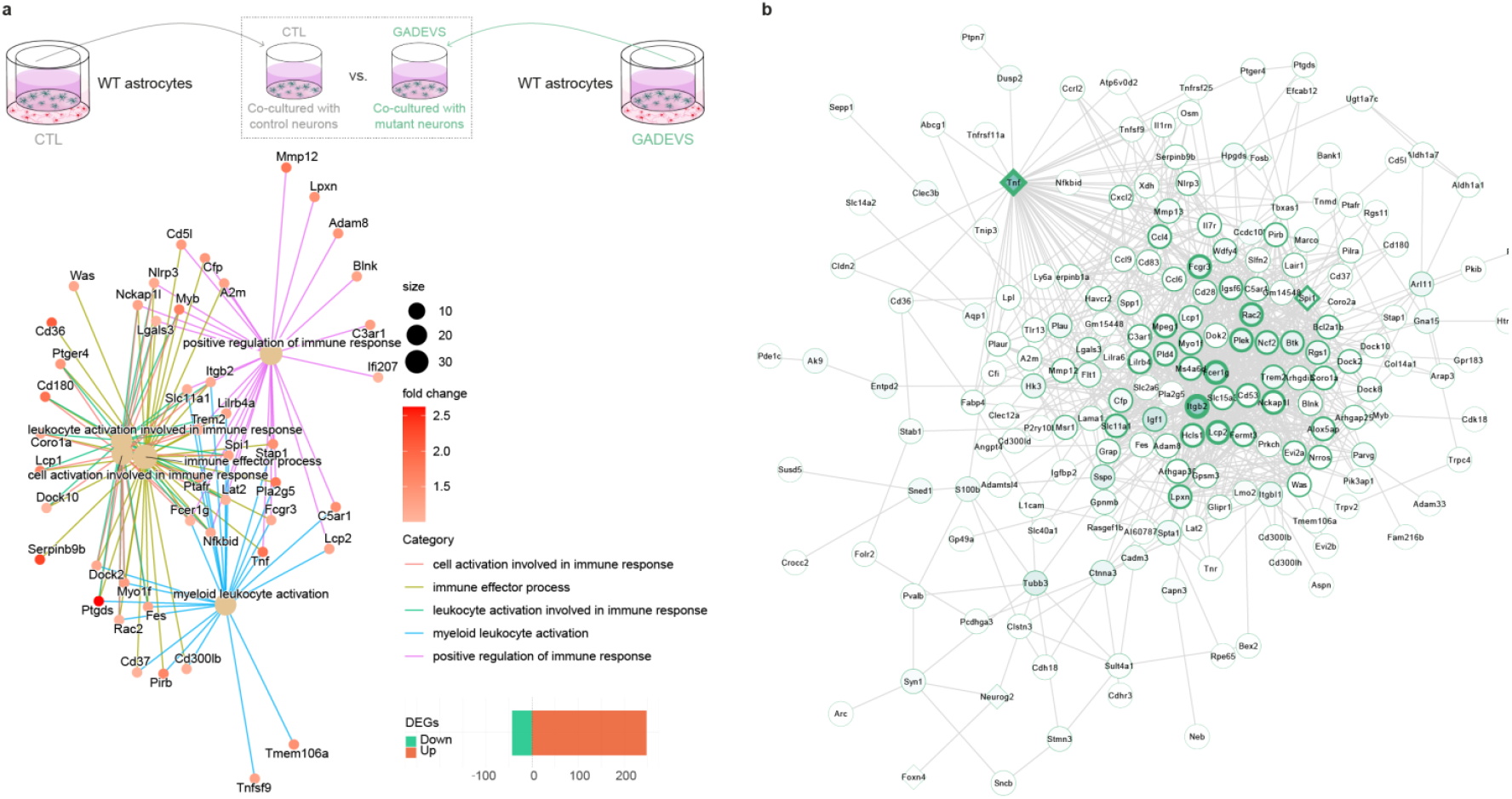
Astrocytes co-cultured with affected neurons undergo cell activation. **a**) Wild type mouse astrocytes co-cultured with both CTL and GADEVS neurons were harvested and profiled using bulk RNAseq. DEA revealed predominant upregulation out of the total 291 DEGs (248 upregulated DEGs, FDR < 0.05 and FC > |2|). Gene-concept network plot describes the interactions across genes belonging to significant biological processes GO terms. **b**) Main component of co-expression network generated from DEGs enriching *cell activation* GO categories depicted in panel **a**. Nodes of TFs are indicated as squares (*Spi1, Myb, Fosb, Foxn4, Neurog2*, and *Tnf*). Edges are denoted as light grey strokes. Node borders are proportional to its degree (highest degree scored 5, DEGs linked are *Itgb2, Tnf, Fcer1g, Lcp2*, and *Fcgr3*). The color of the node span from white (lowest) to green (highest). Nodes with highest centrality scores include *Itgb2, Igf1, Fcerg1, Tnf*, and *Sspo*.

### Mechanistic dissection of YY1-mediated dysregulations in glutamatergic neurons revealed a broad feedback loop attempting at rescuing *YY1* haploinsufficiency and a pervasive role of NEUROG2 in counteracting YY1 activity across biological processes

GADEVS glutamatergic neurons displayed transcriptional and synaptic dysregulations that were sufficient to trigger astrocytic activation, shown by upregulation of gliosis signatures. Thus, to causally pinpoint the disruptions in YY1 regulatory activity, and identify its regulatory partner we leveraged the contextual profiling of transcription and chromatin accessibility in neurons via multiomics. This allowed us to identify differential TF activity and generate cell-type specific GRN to assess the YY1 dysfunctions in mature cortical excitatory neurons.

First, we verified the consistency of the single-cell ATAC modality of NGN2-derived neurons with respect to the RNA modality, by transferring cell-type annotations from the latter onto the UMAP of the former (fig. 7a). Then, we quantified TF differential activity to identify the most disrupted regulators (fig.7b). We identified a general depletion of accessibility for most differentially accessible motifs, and a strong increase in accessibility in fewer TFs, including NEUROG2 and NEUROD1. To reconstruct the contribution of differentially active TFs to the differential expression previously identified in GADEVS mature neurons, we first performed a GO enrichment analysis on the differentially expressed genes, and then measured the enrichments within each enriched GO category for the direct targets of YY1 and those of TFs that simultaneously display significant differential activity and differential expression in neurons (fig.7c). By doing so, we found *ETV5* to be the most strongly enriched for a subset of rather distinguished categories, such as *hypoxia, morphogenesis* and *behavior*, and we found *NEUROG2* enriched in all categories, including *cell adhesion*, and developmental processes. Thus, to further pinpoint the molecular underpinnings of such dysregulations, we extended our gaze to the contribution of these differentially active and differentially expressed TFs to the regulation of all DEGs in mature neurons and plotted the number of shared targets between each TFs and YY1 (fig. 7d). We found a large portion of DEGS to be target of MYC and its two partners MXI1 and EBF3, immediately followed by NEUROG2. Thus, to pinpoint the mechanistic co-dependency between these TFs, we separately derived networks of their reciprocal regulation for control and GADEVS samples and analyzed interactions specific to each genotype to observe potential disruptions in this crosstalk in the patients. In the controls (fig. 7e), we found YY1 as a regulator of NEUROG2, together with KLF5, which appeared involved in a self-repressive interaction with YY1. The GADEVS-specific GRN showed a complete rewiring of this network. Apparently, YY1 haploinsufficiency triggered the contextual down-regulation of *NEUROG2, MXI1*, and *HOXA5*, and the up-regulation of *EGR1, EBF3* and *ETV5*. Concomitantly, the rewired network highlighted a coordinated attempt at upregulating YY1, which included a positive regulation of YY1 by NEUROG2 (fig. 7f).

**Fig. 7.**
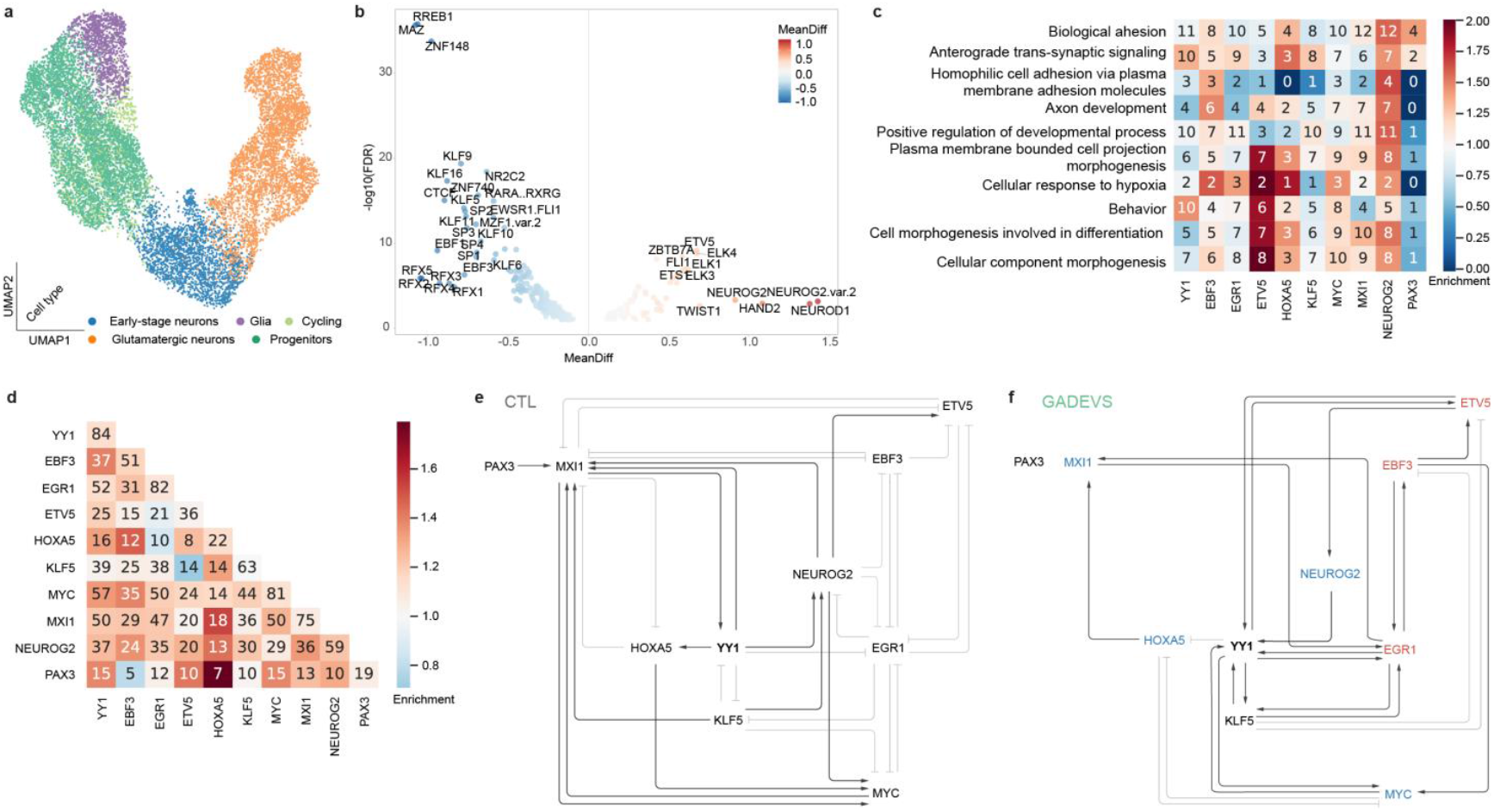
Mechanistic dissection of molecular dysregulations in GADEVS iPSC-derived NGN2 neurons depicts a broad loss of TF activity, and a subnetwork of transcriptional regulation pinpointing YY1 haploinsufficiency to pathologically relevant processes. a) UMAP representation of single-cell ATAC data. Cells are colored by transferring cell-type annotation from each cell shared with the RNA-seq modality. Cell-type specific colors are reported on the bottom of the panel. b) Differential TF activity measured with ChromVar by inference from accessibility changes between control and GADEVS neurons at regulatory regions identified by co-accessibility with Cicero. MeanDiff represent the differential TF activity measured by ChromVar. c) heatmap of GO (biological process) categories enriched by differentially expressed genes. Number of DEGs enriched in each category under the control of each TF (reported below) is indicated in each square. Each square is colored by enrichment measured by the representation of targets of each TF in each category. d) Heatmap of shared targets between pairs of TFs among differentially expressed genes. Each square is coloured by the relative enrichment measured on the targets of the TF indicated on the rows with respect to the target indicated in the column. e-f) Subnetwork of control (e) and GADEVS-specific (f) regulatory interactions among the TFs reported in panels c and d. Arrows and T-shaped edges respectively represent activation and repression (as inferred by CellOracle). Blue and red colored gene names respectively denote downregulation and upregulation of the TFs in GADEVS glutamatergic neurons.

## Discussion

We previously showed that *de novo* mutations of *YY1* cause GADEVS through haploinsufficiency-dependent defects in the activation of YY1-bound enhancers^1^. Consistently, YY1 was found shortly thereafter to act as a general enhancer-promoter looping factor^10,11^. However, genome-wide studies on *YY1* following its acute depletion assayed its role in regulating the 3D genome architecture and transcription initiation levels debated its looping role^14^. Nevertheless, these studies were done in mouse embryonic stem cells, and did not account for the effects of *YY1* in differentiation. Therefore, we hypothesized that YY1 has a role in establishing cell-type identity rather than maintaining pluripotency, altering specific differentiation axes. Consequently, to identify the physio-pathologically relevant cell-type specific impact of *YY1* mutations, we modelled GADEVS neurodevelopment by leveraging patients derived iPSCs, 2D and 3D neuronal lineages. First, in iPSC we identified a broad transcriptional downregulation explained by a global loss of YY1 across different pathogenic truncating or missense mutations. Notably, dysregulated genes in iPSC significantly relate to physio-pathological phenotypes, reinforcing the importance of early developmental settings in revealing the molecular priming of neurodevelopmental conditions^18^.

To define the impact of GADEVS-causing *YY1* mutations on corticogenesis we harnessed patterned cortical brain organoids. Despite an overall similarity in cellular composition in terms of both neuroepithelial and downstream progenitor cells at early developmental stages, we found a striking and highly reproducible faulty organization of ventricle-like structures within GADEVS organoids. Studies conducted across various animal models^8,33,50–54^ had also found evidence of dysfunctional VLS features, hence our observations underscored the fidelity of our model in capturing fundamental aspects of neurodevelopment with the first recapitulation of a pseudo-ventricle phenotype in human organoids which emerges as earliest correlate of the morphological brain phenotype detected in GADEVS individuals with MRI or CT scans^1,15,33,55^. Subsequently, at 30 days of organoid differentiation when neurogenesis kickstarts following the expansion of neuroepithelial cells, with progenitor diversification and the emergence of neuronal lineages, we observed a very specific dysregulation affecting intermediate progenitors. Following on the course of differentiation, late stages (90 days) manifested further major differences in terms of neuronal subtypes among lower and upper layer neurons, with dysregulated YY1 direct targets pointing to disruption of ion channels expression and dysregulation of cell-cell signaling in the EGFR, MAPK, WNT, TNFα and NF-KB (indicative of glial-derived inflammatory processes) pathways.

The cell-cell signaling and synaptic dysregulation were further probed in the 2D NGN2-driven neuronal differentiation paradigm. This revealed in neurons a profound transcriptional rewiring induced by YY1 mutations, subverting the regulons of 6 main transcription factors (*NEUROG2, MXI1, HOXA5, EGR1, EBF3* and *ETV5)* and thereby unmasking a physiological feedback loop between YY1 and NGN2, whose positive regulation by NEUROG2 on YY1 proved however ineffective in rescuing the haploinsufficient setting. Finally, YY1-defective neurons exerted a profound effect on neighboring astrocytes, triggering an inflammatory and astrogliotic response that represents, alongside altered pseudo-ventricles, yet another remarkable endophenotypic correlate of specific alterations observed in the frontal cortex of GADEVS affected individuals^5^. In sum, our work defined the activity of YY1 in human neuronal lineages, across multiple stages of development, highlighting both YY1 regulatory activity and its downstream effects on neuronal development and mature neuronal functions, and extending the understanding of YY1 haploinsufficiency to non-cell autonomous alterations. As with any condition of such complexity, this multiscale dissection of endophenotypes can now guide further modelling studies with the aim of identifying the most actionable targets that relay the molecular mechanisms exposed by single cell multiomics to the main cell biological defects at the level of pseudo-ventricles, synaptic puncta and reactive astrocytes. In this regard, given that even astrocyte malfunctions alone are sufficient to induce changes in basal transmission and plasticity processes^61,^ our findings of neuronal-mediated inflammation points to look at inflammation modulation as a potential path ahead for preclinical testing in GADEVS.

As an increasingly recognized condition^6,33^ GADEVS calls for a matched research and care strategy aimed at ever more accurate screening and detailed phenotyping of case reports to refine clinical description and align it to the underlying molecular mechanisms. Therefore, we established a website to collect detailed phenotypic data of individuals harboring YY1 mutations not only to gain insight into the clinical spectrum that these mutations might cause but also to obtain fundamental understanding of the pathogenic mechanisms underlying the Gabriele-de Vries syndrome (https://humandiseasegenes.nl/yy1). With this first modelling of GADEVS, we provide a high-resolution definition of its key pathogenic mechanisms, uncovering multiscale longitudinal phenotypes, cell-autonomous and non-cell autonomous alike, whose multiomic underpinnings and functional readouts orient the search for druggable targets and repurposed compounds in physio-pathologically relevant human neuronal lineages ^60–62^.

## Materials and Methods

### Cell reprogramming

Human fibroblasts were isolated from skin biopsies and lymphoblastoid cell lines (LCLs) were generated by Epstein-Barr virus (EBV) transformation of the B-lymphocytes within the peripheral blood lymphocyte leading to proliferation and subsequent immortalization of these cells. Both healthy and non-healthy human samples (fibroblasts and LCLs) were reprogrammed using the non-integrating Sendai virus (SeV) method to safely and efficiently deliver and express key genetic factors (Yamanaka factors OCT3/4, SOX2, KLF4, and c-MYC). For all lines, a total of 3 × 10^5^ cells were transduced per individual using a multiplicity of infection (MOI) of 5-5-3 (KOS comprising human Klf4, Oct3/4, and Sox2; c-MYC containing human c-Myc; Klf4 with human Klf4, respectively). Moreover, the amount of virus used was calculated according to each certificate of analysis respective of each SeV kit. The reprogramming procedure was performed according to the manufacturer’s protocol using feeder-dependent protocols for both fibroblasts and PBMCs (feeder layer containing mouse embryonic fibroblasts). Cells were maintained every day and small colonies were subsequently picked and expanded approximately 1-1.5 months post-transduction. After reprogramming from LCL, EBV expression in iPSCs was evaluated, and samples completely turned off EBV expression compared to a positive control cDNA taken from LCL, showing irrelevant Ct, and with melting curves comparable with the negative (no cDNA). Reprogrammed fibroblast-derived iPSCs were also validated to be free of SeV transduced vectors by RT-qPCR. iPS cells were pulled down and pelleted for RNA extraction and depletion was checked by RT-qPCR according to manufacturer’s instructions. A reverse transcription was carried out using 2.5 μg of RNA following the instructions provided with the kit (Cat Nº, 11766050, Invitrogen). For PCR reaction mix preparation, 12 ng of cDNA for each sample was used and performed in triplicates using the Fast SYBR Green Master Mix (visualize primers in table S1). Positive control cells were set aside during the first stages of the reprogramming procedures (fig. S1, Cat Nº A16517, ThermoFisher).

### Cell culture

#### Fibroblast/LCL handling and iPSC culturing

Human fibroblasts were cultured in DMEM, FBS 10%, non-essential amino acids (NEAA) 1%, penicillin-streptomycin (P/S) 1%, and β-mercaptoethanol 0.1%. LCLs were grown in RPMI 1640, FBS 15%, HEPES 1%, L-glutamine (Gln) 1%, and P/S 1%. Trypsin was used to passage fibroblasts whereas LCLs were cultivated in suspension and expanded by dilution. Human iPSCs were cultured in feeder-free conditions onto matrigel-coated dishes using TeSR-E8 medium supplemented with P/S 1% and passaged using either ReLeSR (Cat Nº 100-0483 stemcell technologies, generates cell aggregates for expansion and standard maintenance) or accutase solution (Cat Nº A6964 Sigma, exclusively for experimental procedures) supplemented with ROCK inhibitor 5 μM (Y-27632 dihydrochloride, #A3008 ApexBio) was added to the culture to enhance single cell survival. Cryopreservation of hiPSCs was performed in complete TeSR-E8 medium with 10% DMSO and supplemented with ROCK inhibitor 5 μM. All samples were routinely tested for mycoplasma.

#### Directed differentiation of hiPSCs into ectoderm, mesoderm, and endoderm lineages

To functionally validate the potential of human iPSC lines to differentiate into all three embryonic germ layers, we took advantage of the STEMdiff™ trilineage differentiation Kit and followed the manufacturer’s instructions with minor modifications (Cat Nº 05230, Stemcell technologies). hiPSCs were cultured onto Matrigel-coated glass coverslips and on day 0, 2.5 × 10^5^, 7.5 × 10^4^ and 2.5 × 10^5^ cells were plated for further differentiation into ecto-, meso-, and endoderm, respectively. At the end of differentiation, cells were fixed, and lineage-specific markers were used for immunofluorescence using the following established markers NESTIN and PAX6 for ectoderm; Brachyury and CXCR4 for mesoderm; and FOXA2 and SOX17 for endoderm (fig. S2).

#### Differentiation of iPSCs into neural crest stem cells (NCSCs)

hiPSCs were differentiated into NCSCs as previously described^63^. NCSC differentiation required 15-20 days and was carried out as follows: 90% confluent iPSCs were detached with accutase solution and plated onto matrigel-coated dishes using TeSR-E8 medium supplemented with 5 μM ROCK inhibitor at a density of approximately 9.2 × 10^4^ cells per cm^2^. After 24h, NCSC differentiation medium was added and changed every day. NCSC medium was composed of DMEM-F-12 1:1, 10% probumin (stock solution of 20% m/v in DMEM F-12 1:1), P/S 1%, L-Glutamine 1%, NEAA 1%, trace elements complex 0.1%, 0.2% β-mercaptoethanol (50 mM), Transferrin (10 μg/mL), sodium L-ascorbate (50 μg/mL), Heregulin-1 (10 ng/mL), LONG®R3 IGF-I (200 ng/mL), FGF2 (8 ng/mL), GSK3 inhibitor IX Bio (3 μM) and SB431542 (20 μM). Cells were passaged every 4-5 days and plated at high densities. Upon differentiation, NCSC were stocked as stable lines and cultured using NCSC medium with routine splitting ratios of 1:4-5 (fig. S3e,f).

#### Preparation of primary mouse astrocytes

Primary astrocyte cultures were created from the cerebral cortices of embryonic day 18 (E18) mice embryos and maintained as previously described^56,57^. In a nutshell, after embryo collection, cortices were dissected from each embryo’s brain, and chemical dissociation with trypsin (2.5%, at 37 °C for 30 min) was performed. Afterwards, mechanical dissociation was performed by vigorous pipetting. Following digestion, cortex tissues were centrifuged (300 g for 5 min) and supernatant discarded. Finally, astrocyte medium (DMEM supplemented with DMEM medium supplemented with FBS 20%, P/S 1%) was added and 1-1.5 × 10^7^ cells were seeded in each T75 culture flask. After reaching confluence (10-15 days), astrocytes were passaged with trypsin and expanded. This protocol allowed the generation of pure astrocytic cultures due to the gradual absence of viable neurons and other glial cells. Astrocytes were generated and maintained 3-4 weeks prior NGN2-derived neuron differentiation.

#### Differentiation of iPSCs into NGN2-derived cortical neurons

hiPSCs were engineered using a ePiggyBac (ePB) transposon previously electroporated using the Neon™ Transfection System (MPK5000, ThermoFisher). Electroporation parameters for each reaction of 4 × 10^5^ cells were: 900V, 20 ms, 2 pulses using 5 μg of total DNA in a 1:10 ratio of helper:vector (0.5 μg of an helper plasmid expressing a transposase and 4.5 μg of donor plasmid with a transposable element). NGN2 ePB donor plasmid has the following genetic configuration: hUbC promoter - rtTA - T2A - BsdR - TRE - Ngn2 - P2A - EGFP - T2A – PuroR and allows the selection with blasticidin (fig. S7a). Following transfection, cells were allowed to recover for 48h and selection was performed using blasticidin 5 μg/mL until all cells were eliminated from the negative control performed without the addition of the transposase (usually 5-7 days of selection). Stable iPSCs containing the ePB transposon were stored until needed for neuronal differentiation.

Neuronal differentiation is induced by doxycycline and is followed by NGN2 overexpression, as previously described^64^. Cortical neurons were maintained in Neurobasal medium fully supplemented (B27 with vit.A + P/S + Glutamax + doxycycline + puromycin + N2 + NEAA + human Laminin + NT3 + BDNF). Fresh mouse astrocytes were added to the culture by means of hanging cell culture inserts (MCRP06H48, Millipore, transwell for clarity) on day 4 after definitive plating of neurons in poly-d-lysine-coated 6 well plates or added into poly-L-ornithine-coated glass coverslips treated with nitric acid for IF experiments at a ratio 1:1 (human neuron to mouse astrocyte).

#### Differentiation of iPSCs into 3D human *in vitro* systems – cortical brain organoids

Cortical brain organoids were generated using an adaptation of a previously described protocol^36^. Prior to organoid culture, iPSCs were instead routinely cultured with mTeSR™1 basal medium (Cat Nº 85850 stemcell technologies). hiPSC were expanded in 10 cm dishes and detached at 60-70% confluency with Accutase for obtaining single cell suspensions. After centrifugation (150 g, 3 min), cells were resuspended in mTeSR™1 basal medium supplemented with ROCK inhibitor (5 μM), TGF-β inhibitor SB-431542 (10 μM), and dosromorphin (1 μM) and counted.

To form free-floating spheres, approximately 4 × 10^6^ cells were added to a single 6-well plate and kept under rotation (95 rpm) inside the incubator (37 °C, high O_2_ conditions) for 24 h. The medium was refreshed every day by removing half (2 mL) and adding half (2 mL) for the following 2 days. After 3 days (day 4 to 10), supplemented mTeSR™1 basal medium was replaced by Media 1, comprised of Neurobasal A medium supplemented with 1% GlutaMAX, 1% N2 NeuroPlex, 1% B27 with vitamin A, 1% NEAA, 1% P/S, SB (10 μM) and Dorsomorphin (1 μM). Subsequently, from days 11 to 17, cells were maintained in Media 2: Neurobasal A medium with GlutaMAX, 1% B27 + vitamin A, 1% NEAA and 1% P/S supplemented with FGF2 (20 ng/mL), followed by 7 additional days (day 18 to 24) in Media 2 supplemented with FGF2 (20 ng/mL) and EGF (20 ng/mL) to boost neural progenitor proliferation. Finally, organoid medium conditions shifted to Media 3, composed of Media 2 supplemented with BDNF (10 ng/mL), GDNF (10 ng/mL), NT-3 (10 ng/mL), L-ascorbic acid (200 μM) and dibutyryl-cAMP (1 mM) to promote network maturation, gliogenesis and enable neuronal activity. After 7 days (31 days *in vitro*), cortical organoids were maintained in Media 3 for as long as needed, with media changes every 3-4 days. Catalog numbers for all reagents and small molecules used for the cell culture medium were previously described^36^.

### RNA extraction and library preparation for bulk RNA sequencing (RNAseq)

Cultured iPSCs and NGN2-derived neurons were washed with phosphate-buffered saline (PSB) and treated with 600 μL RTL buffer supplemented with 10% β-mercaptoethanol. Afterwards, total RNA was extracted using the RNeasy Mini Kit. Purified RNA was quantified using a NanoDrop spectrophotometer and RNA quality was further checked with an Agilent 2100 Bioanalyzer using the RNA nano kit (Cat Nº 5067-1511, Agilent). Only samples with RIN>9 were used for library preparation. Prior to library preparation, ERCC spike-in mixes (Cat Nº 4456739, ThermoFisher) were added to all samples to facilitate data normalization and quantification. RNA sequencing libraries were prepared following manufacturer’s protocols for Truseq-stranded Total RNA (RiboZero depletion) or using Illumina Stranded mRNA Prep, starting from 250-500 ng of total RNA. cDNA library quality was assessed on Agilent 2100 Bioanalyzer, using the Agilent High Sensitivity DNA Kit (Cat Nº 5067-4626, Agilent). Libraries were then sequenced with the Illumina Novaseq 6000 instrument at a read length of 50 bp (paired-end) with a coverage of 35 million reads per sample.

### Chromatin immunoprecipitation followed by sequencing (ChIP-Seq)

ChIP-seq for TF YY1 was performed in all iPSC lines (4 CTLs *vs*. 3 GADEVS lines) using approximately 1 × 10^7^ cells per sample. First, cells were harvested and crosslinked with formaldehyde 1% in PBS for 8 minutes at RT under continuous rotation. To quench the reaction, glycine (125 μM) was added to each mix and incubated for 5 min on ice. Cells were subsequently pelleted by centrifugation (500 g, 3 min at 4 °C) and washed in ice-cold PBS. Cells were lysed in 10 mL buffer A (50 mM HEPES pH 8.0, 140 mM NaCl, 1 mM EDTA, 10% glycerol, 0.5% NP-40, 0.25% Triton X-100) for 10 minutes on ice. After centrifugation, cell pellet was resuspended in 5 mL buffer B (10 mM Tris pH 8, 1 mM EDTA, 0.5 mM EGTA and 200 mM NaCl) and incubated for 5 min on ice. Next, nuclei were pelleted (500 g, 3 min at 4°C), resuspended in 150 μL buffer C (50 mM Tris pH 8, 5 mM EDTA, 1% SDS, 100 mM NaCl, 1x Roche complete mini protease inhibitors) and incubated on ice for 10 min. Afterwards, 350 μL ice-cold TE buffer was added and chromatin was sheared in 1.5 mL microtubes using a Bioruptor Pico sonication device (B01060010, Diagenode) for 3 × 5 cycles (30 s ON/ 30 s OFF) at 4°C. Upon addition of 55 μL of 10x ChIP buffer (0.1% SDS, 10% Triton X-100, 12 mM EDTA, 167 mM Tris-HCl pH 8, 1.67 M NaCl), chromatin was centrifuged (13 000 g, 10 min at 4 °C). At this step, 1% of sheared chromatin was saved as input control, whereas the rest was transferred into fresh tubes. 10 μg of antibody was added for YY1, and 5 μg for normal rabbit IgG immunoprecipitations (antibody information on table S2) and were then incubated overnight on a rotating wheel at 4 °C. The following day, 40 μL of protein G Dynabeads (mixed 1:1 and washed with 1x ChIP buffer) were added and incubation was continued for another 4 h. Each ChIP were washed: 2 × LSB (10 mM Tris-HCl pH 8.0, 1 mM EDTA pH 8.0, 140 mM NaCl, 1% Triton X-100, 0.1% SDS, 0.1% Na-deoxycholate); 3 × HSB (10 mM Tris-HCl pH 8.0, 1 mM EDTA pH 8.0, 360 mM NaCl, 1% Triton X-100, 0.1% SDS, 0.1% Na-deoxycholate); 1 × LiSB (10mM Tris-HCl pH 8.0, 1mM EDTA, pH 8.0, 250mM LiCl, 0.5% NP-40, 0.5% Na-deoxycholate); and 1 × TE (10mM Tris-HCl pH 8.0, 1mM EDTA, 50mM NaCl).

Beads were transferred into a new tube during the last wash, and wash buffer was completely removed before adding 100 μL of elution buffer (10 mM Tris-HCl pH 8.0, 1 mM EDTA pH 8.0, 150 mM NaCl, 1% SDS). Mix was incubated for 30 minutes at 65°C with constant shaking, and after 2 μL RNaseA (20 μg/ μL) were added, samples were incubated for 1 h at 37°C. Input samples were adjusted to 150 μL total volume with elution buffer and processed similar to ChIP samples. Then 2 μL Proteinase K (20 mg/mL) was added and samples were incubated 2 h at 55°C and then O/N at 65°C. After two volumes of AMPure XP beads and one volume of Isopropanol were added, samples were vigorously mixed and incubated for 10 min at RT. Next, beads were collected on a magnetic rack, washed twice with 80% EtOH, and DNA was finaly eluted in 40 μL 10 mM Tris pH8.0 for 5 min at 37°C. DNA libraries were prepared and sequenced on the Illumina Novaseq 6000 instrument at 50 bp paired-end read length and a coverage of 35 million reads per sample.

### Single cell multiomics (gene expression + chromatin accessibility)

NGN2-derived neurons were collected, and single cell suspension was immediately processed. As opposed to neurons, cortical organoid dissociation required a more tailored and gentle procedure. 30- and 90-days cortical organoids were dissociated using a papain-based dissociation buffer composed of 30 units/mL papain (Cat Nº 07466, stemcell technologies), dissolved in filter-sterilized activation solution comprised of 1.1 mM EDTA, 0.067 mM mercaptoethanol, 5.5 mM L-cysteine HCl) and 125 units/mL DNase I (07900, Stemcell technologies) in Hanks’ Balanced Salt Solution (HBSS) with 10 mM HEPES, without phenol red (Cat Nº 37150, stemcell technologies). Pools of homogeneously sized organoids (4-8 organoids depending on timepoint) were incubated on a rotating wheel (approximately 30 min at 37 °C) with 1 mL of activated dissociation buffer, followed by manually pipetting. Afterwards, cells were transferred into a new eppendorf leaving behind undissociated pieces, and further centrifuged (300 g, 5 min at 4 °C). Supernatant was removed and cells were resuspended in 1 mL of PBS-BSA 0.04% and filtered with 40 μm Flowmi cell strainer to remove residual aggregates and debris. Single cell suspensions were counted manually (1:1, single cells, and trypan blue), and different lines were multiplexed together to obtain a final number of 1 million cells, equally mixed (e.g., 250k cells in case of 4 samples multiplexed), assembling a single reaction. This step allowed to pool together different genotypes which can later be de-multiplexed based on their transcriptome previously profiled with bulk RNAseq (at iPSC stage or NGN2-derived neurons).

Nuclei Isolation for single cell multiome (ATAC + Gene expression sequencing) was performed according to the manufacturer instructions in the demonstrated protocol from 10X Genomics (CG000365-RevB). Briefly, each reaction was washed twice with PBS-BSA 0.04%, and pellet was resuspended in 100 μL of lysis buffer (10 mM Tris-HCl pH 7.4, 10 mM NaCl, 3 mM MgCl2, 0.1% Tween-20, 0.1% Nonidet P40 Substitute, 0.01% digitonin, 1% BSA, 1 mM DTT, 1 U/μL RNase inhibitor, nuclease-free water), and incubated for 3 min on ice. Lysis buffer was washed away three times with 1 mL of wash buffer (10 mM Tris-HCl pH 7.4, 10 mM NaCl, 3 mM MgCl2, 1% BSA, 0.1% Tween-20, 1 mM DTT, 1 U/μL RNase inhibitor, nuclease-free water). Assuming 50% nuclei loss during lysis, nuclei were resuspended in diluted nuclei buffer (1X Genomics Nuclei Buffer, 1 mM DTT, U/μL RNase inhibitor, nuclease-free water) according to the reference table provided by 10X protocol appendix. The volume of nuclei diluted buffer is critical to fit the right range of concentration based on the number of targeted nuclei recovery, therefore avoiding overcrowding of the Chromium machine during tagmentation and GEM preparation steps. Resuspended nuclei were passed again through a Flowmi cell strainer and counted to fit the right concentration for the number of targeted nuclei (5 000 was the number of targeted nuclei recovery for each multiplexed sample in all the experiments). DNA libraries were prepared according to the manufacturer’s recommendations (10X Genomics) and further sequenced on the Illumina Novaseq 6000 instrument at a coverage of 50 000 reads per nucleus.

### Western Blot

Human iPSCs were grown as previously described, pelleted (300 g, 5 min), and then resuspended in 100 μL RIPA buffer (50 mM Tris-HCl, pH 7.5, 150 mM NaCl, 1% Triton X-100, 0.5 mM EDTA, and 5% glycerol) supplemented with protease inhibitor cocktail (PIC) and 1 mM PMSF. Total protein lysate concentration was measured using the Biorad protein assay. For western blotting, 30 μg of protein were resolved on polyacrylamide gels (12% separating gel), which were transferred on nitrocellulose membrane. Blocking was performed for 45 min in 5% non-fat dry milk in TBS plus 0.05% Tween 20 (TBST), and membranes were incubated with primary (overnight at 4 °C) and secondary (2 h at RT) antibodies. Signal was detected with corresponding horseradish peroxidase (HRP)-conjugated secondary antibodies (antibody information on table S4), imaged with ChemiDoc XRS + System (Biorad), and quantified.

### Immunofluorescence, image acquisition, and data analysis

#### Immunocytochemistry using 2D samples

Control and mutated cells (iPSCs, human-derived neurons, and mouse astrocytes) were cultured onto nitric acid-treated glass coverslips as previously described until wash (PBS) and fixed (4% methanol-free paraformaldehyde for 10 min at RT). For immunofluorescence (IF), fixed cells were then permeabilized with 0.2% Triton X-100 diluted in PBS for 10 min and blocked for 1 h in 5% serum matched with the species of the secondary antibody (donkey serum). Next, samples were incubated with the primary (overnight at 4°C, antibodies described in table S2) and secondary (1 h at RT) antibodies diluted in blocking buffer. Antibodies were removed by washing with PBS in between incubations. Afterwards, cells were stained with DAPI (1:5 000) for 5 min, washed with PBS, and mounted onto glass slides with Mowiol-Dabco mounting medium. Samples were stored at - 20 °C until visualized in the confocal microscope.

#### Acquisition and data analysis for counting synaptic puncta

Following fixation, staining, and mounting into glass slides, human-derived neurons were visualized using a Leica SP8 Confocal microscope equipped with acousto-optical beam splitter (AOBS), resonant scanner, motorized stage x-y-z as well as DM camera for widefield acquisition (Leica Microsystems, Germany). Previews of the whole coverslip were taken in widefield mode (10X/0.3 dry) using the DAPI channel to choose continuously the central area of the slide, that was further acquired at a higher resolution in confocal mode using the LASX software equipped with navigator (version 3.1.5.16308). Three-channel (MAP2, Synapsin and DAPI) z-stack images (z-step intervals of 1 μm) were sequentially acquired using a 63X/1.4 oil objective and a DFC365 FX CCD Camera (Leica) with a x-y sampling of 72 nm. A total of 10 fields of view were acquired for each glass slide (with 4 slides being imaged, 40 fields of view per sample imaged and analyzed). After acquisition, each field of view was analyzed individually using an open-source software followed by custom design of a semi-automated macro (FIJI-ImageJ v2.1.0, USA, macro 1) by segmentation the C1 (MAP2B) and C2 (Synapsin1/2) channels for further analysis. First, all C1 channel fields of view were transformed into binary images and converted to masks, from which the area was measured to serve as normalizer as well as used to capture exclusively synapsin1/2-positive signal co-localized within MAP2B-positive structures. Then, synapsin1/2-positive particles were measured, ROIs saved, and counts normalized in relation to MAP2B area for each field of view. All data is presented as median of *n* different fields of view for all conditions (transformed as percentages).

#### Immunostaining and clearing of 3D structures - cortical brain organoids

Whole cortical organoids were immunostained and cleared using the MACS^®^ Clearing Kit (Miltenyi Biotec, 130-126-719) and following entirely the manufacturer’s instructions (Immunostaining and clearing of organoids and spheroids for 3D imaging analysis) comprising some minor modifications. Organoids were collected at days 12, 30, and 90 and were washed in PBS followed by fixation using 4% methanol-free paraformaldehyde (20 min at RT). Following multiple washes to remove the fixative agent, organoids were permeabilized (6 h at RT) under continuous rotation. Incubation with primary antibodies was perfumed for 2 days at 37 °C under rotation (antibodies listed in table S2). To remove unbound antibodies, antibody staining solution was discarded and washed 5 times for 30 min, with slow continuous rotation. Secondary antibodies were diluted according to manufacturer’s recommendations and added to each well (incubated for 1 h at RT). If required, DAPI staining was performed at this step (10 min at RT). Afterwards, antibody staining solution was discarded and replaced with fresh ones (30 min incubations at RT). These steps were repeated 5 times to ensure full removal of unbound antibodies. Organoids were afterwards embedded in 1% agarose in ddH_2_O and following solidification were cut into approximately 2×2-5×5 mm pieces (depending on organoid size), each containing one or more organoids. Smaller agarose cubes facilitate imaging conditions, and many orientations ensure visualization of the entire organoid. Dehydration solutions were prepared by diluting absolute ethanol in sterile water to obtain 50% and 70% ethanol solutions. Up to ten embedded organoids were dehydrated with a series of ethanol dilutions in 1.5 mL tubes at RT under slow continuous rotation: 50% ethanol was incubated for 2 h, followed by 70% ethanol for 2 hours and finally 100% ethanol overnight. The next day, clearing solution was added into 1.5 mL tubes containing dehydrated organoids, and incubated at RT under slow continuous rotation for a total of 6 h. Clearing Solution was discarded and substituted with imaging solution to proceed with imaging acquisition and long-term storage (protected from light).

#### Sample acquisition and data analysis

Whole brain organoids were further visualized in a Yokogawa Spinning Disk Field Scanning Confocal System (Yokogawa CSU-W1 25 μm - 50 μm pinhole dual disk, Nikon, Japan), equipped with motorized stage x-y-z, and a Prime BSI camera (Teledyne Photometrics, Arizona, USA), in confocal mode. Four channel (DAPI, NESTIN, PAX6, and SOX2) z-stack images of the whole organoid (z-step intervals of 5 μm) were acquired using the Nikon NIS Elements AR software (version 5.02.03) at a 10x/0.3 magnification (dry, no binning). For counting cells actively undergoing mitosis in the two conditions, images were processed in the Arivis Vision 4D software (version 3.5.0, Arivis, Germany), using a fully automated custom pipeline. Briefly, to facilitate the downstream 3D workflow, DAPI and pHH3 channels were processed separately, and advanced image enhancement filters were applied. For volume measurements, the DAPI channel in each organoid was later segmented using the “Li” thresholder, and objects smaller than 10 000 μm^3^ were filtered out. For counting the number of pHH3-positive cells, the “Blob Finder” analysis operator was used to segment cells and the following parameters were applied: i) averaged diameter of 6 μm; ii) 5% probability threshold; iii) 60% split sensitivity. Finally, raw data was processed, and the number of particles was normalized to DAPI volume. All data is presented as median of *n* different organoids for all analysis. For analysis of cytoarchitectures and classification of ventricle-like structures (VLS) in CTL- and GADEVS-derived organoids, VLS measurements were classifies as “regular” or “irregular”. Given the empirical/arbitrary logic of the analysis, in order to ensure bias-free annotation of “regular” and “irregular” VLS, individual files containing a single organoid were anonymized and scrambled (named with 5 random characters) before manual annotation. Each structure was eligible for classification in “regular VLS” if the following criteria were met: i) structure is PAX6-positive; ii) structure has clear PAX6-lined ventricle lumens; iii) structures were sufficiently separated from neighboring structures to ensure accurate counting; iv) truncated structures where we could not discern between start and end of VLS were not counted (verified with 3D rendering of each organoid). For classification of structures in “irregular VLS”, the following premises include: i) structure is PAX6-positive; ii) structure lack well-defined lumen; iii) structures were adequately apart, otherwise were not counted.

### Morphometrical characterization of organoids

Organoid morphometric properties were assessed for each individual cell line and bright-field images were acquired using the OLYMPUS IX81-ZDC inverted microscope, equipped with a Hamamatsu ORCA-ER B/W CCD camera at a magnification of 4X/0.16 (dry) without binning. Image acquisition was fully automated, and analysis was performed using Fiji software (FIJI-ImageJ v2.1.0, USA). Two complementary macros were developed to first stitch multiple fields of view (see Macro 2) and afterwards perform the morphological analysis (see Macro 3). Two distinct methods were used to detect objects according to organoid density for each dataset. When handling crowded acquisitions, the plugin ‘StarDist’ was used to automatically select each organoid. Otherwise, samples were segmented and converted into masks and objects were marked for analysis. Regardless of the selection method used, objects touching the edges in each field of view were discarded and analysis followed the same principles in which: i) objects smaller than 25 000 μm^2^ were discarded from the pipeline; ii) contrast was enhanced in all fields of view; iii) images were segmented and binarized; iv) area, perimeter, and circularity were analyzed using Fiji’s ‘Analyze Particle’ plugin; v) data was saved on the region of interest (ROI) manager. The measured parameters correspond to the largest cross-section of each organoid.

### Computational analysis

#### Bulk RNA-seq analysis

All bulk RNAseq data were quantified by means of Salmon v1.4^65^, using reference GENCODE genome hg38 v35 (CRCh38.p13) or GENCODE assembly GRCm39 (release M27), if handling human or mouse datasets, respectively. We derived HUGO Gene Nomenclature Committee (HGNC) symbol gene names from the corresponding general transcription factors (GTF). Each bulk dataset was filtered using a pre-normalization threshold: only genes with at least 20 raw read counts in at least two biological replicates were kept. The same thresholds were used to filter pseudo-bulks, which were generated via adpbulk (*github*.*com/noamteyssier/adpbulk*), aggregating raw reads by cell-type and by individual.

Differential expression analysis was performed using previously published “edg2” wrapper^66^ for bulk RNA-seq and “edg1” for pseudo-bulk, respectively using *estimateGLMRobustDisp* or *estimateDisp* for dispersion estimation. FDR < 0.05 and FC >= 1.25 were used as thresholds in bulk, and FDR < 0.01 and FC > 2 were used in pseudo-bulks. Gene Ontology (GO) enrichments were performed with topGO^67^ and represented with internal scripts on RStudio (R) 4.1.0. enrichGO^68^ was used on genes passing false discovery rate (FDR) 0.05 to generate dotplots and GO semantic networks. Transcription factor enrichments (also “master regulatory analyses” elsewhere) were performed by means of internal R scripts, based on i) recursive hypergeometric tests performed at gene level, by comparing known targets of TF that have been derived from motifs and ENCODE ChIPseq data using TFBS database^69^, with differentially expressed genes (DEGs), using as background universe the expressed genes in the filtered count matrix; ii) enrichment was calculated as follows: observations are set as the intersection of DEGs and targets of each TF; expected observations are measured as the multiplication of lengths of the two genesets, divided by the length of the universe; iii) FDR is measured by Benjamini & Hochberg multiple-test correction.

#### HPO term enrichments were calculated with a hypergeometric test, using all expressed genes as universe. The ggraph R package (v.2.0.6) was used for visualization of the results

Astrocytes network deconvolution was performed as follows: starting from differential expression, we inferred transcription factor activity by means of viper using the DoRothEA database^70^. We performed network analysis in Cytoscape^71^ calculate topological indexes of each node (i.e., DEG), such as degree (calculated as a sum of the weight of all connections of a given node), betweenness (the quantification of the number of times a node is a bridge along the shortest path between other two nodes), and betweenness centrality, which were used as centrality indices. Fosb, Foxn4, Myb, Neurog2, Spi1 and Tnf were all differentially expressed transcription factor with high betweenness and centrality scores, and they were thus predicted to be responsible for differential expression patterns (i.e., master regulators) identified in astrocytes grown with patient-specific neurons.

#### ChIP-seq analysis

Chromatin Immunoprecipitation sequencing (ChIP-seq) reads were trimmed for library specific adaptor contamination before being aligned to the hg38 genome with Bowtie 2 (removing multi-mapping reads with samtools). We performed peak calling via MACS 2.1 using narrow settings for YY1 (-q 0.05). Peaks overlaps were performed by means of bedtools v2.28, and bound genes were defined as intersecting promoters (from 500bp upstream to 250bp downstream TSS), or intersecting enhancers (cell-type specific 4D Genome consortium peak sets). Presence of peaks in exclusion lists from ENCODE was verified. Motif enrichments were performed with Homer. To perform motif enrichment analyses on bulk TF ChIPseq we started from summit peaks, extended them by 100 bp on both sides of the summit using BedTools, intersected all control replicates and kept peaks found in at least 2 of them. Hg38 was used as reference genome and Homer parameters --mask and -size 200 were applied.

Tracks and Heatmaps were generated with Deeptools v3.5^72^. Reference peaks were defined as present in at least 2 control samples. We identified “lost peaks” as regions preferentially found in CTLs (i.e., regions in at least n=1 controls and none of patients, plus regions found in all controls and at most 1 patient). Peaks were considered “gained” following the inverse logic, as regions preferentially found in patient. Motif enrichments were performed with Homer v.4.11 using default parameters. Coverage heatmaps were generated using deepTools plotHeatmap onbamCoverage (RPGC normalization) or bamCompare (normalization on Inputs) ouputs, as stated in the text.

#### Multiomic data preprocessing

Multiomic sequencing data was preprocessed using 10X CellRanger ARC [10.21105/joss.03021] and computationally demultiplexed based on individual-specific single nucleotide polymorphisms using SCanSNP^73^. The scRNA-seq data was analyzed using the Scanpy package ^74^, v.1.7.2). The Matplotlib^75^, v. 3.4.145) and Seaborn^76^, v0.11.146) packages were used for visualization. Pandas (^77^,v1.2.4) and Numpy (^78^, v1.22.3) were used for data handling.

Differential expression analysis between GADEVS and control cells in each cluster was performed by pseudo-bulk. Cells were grouped by cell line and subsampled to have a similar number of cells in each group before summing the raw counts. EdgR (v.3.32.1, ^79^) was then used to perform differential expression between the aggregated counts of the controls and the disease patients: first we normalized by trimmed mean of M values (TMM) using the CalcNormFactors function, then we obtained negative binomial (NB) dispersion estimates using estimateGLMRobustDisp, we applied a GLM likelihood ratio test (glmFit) and found genes differentially expressed between conditions (glmLRT, topTags). Sex was included as a covariate in the design matrix.

The topGO R package (v.2.42.0) was then used for Gene Ontology^80^ term enrichment analysis. We tested for Biological Process, Molecular Function and Cellular Component Terms, using as background all genes expressed in our dataset. We applied a Benjamini-Hochberg p-value adjustment procedure and set a significance threshold of 0.1 on the FDR.

#### scATAC-seq data analysis

The ArchR (^81^, v.1.0.2) R toolkit was used for the main steps of scATAC-seq data analysis. ggplot2 (^82^, v.3.3.4) was used for visualization.

Fragments were filtered by length and counted in genome-wide 500-bp bins. After generating Arrow files and filtering cells based on scATAC-seq-specific quality metrics (see provided scripts), we intersected the cell barcodes with those that passed scRNA-seq QC. Normalization and dimensionality reduction were performed using Latent Semantic Indexing. A UMAP projection was computed for visualization.

After confirming that cell types annotated in scRNA-seq clustered coherently on the scATAC-seq UMAP, the annotation was given as grouping, combined with sequencing batches, to run ArchR’s iterative peak calling using MACS2 (^83^, v.2.2.7). Normalization and dimensionality reduction steps were then recalculated on the resulting peak count matrix. We downloaded motif position frequency matrices from the JASPAR database (2020 version,^84^) and added motif information to the ArchR object. We used ChromVar (v. 1.16.0, ^85^) to compute per-cell motif activity scores. We then tested for differential TF activity scores between GADEVS and control samples by applying a Wilcoxon Rank Sum Test and computing the average difference in z-score between the two groups.

#### Promoter-enhancer association

We used Cicero (^86^, v. 1.8.1) to find cell type-and condition-specific co-accessible peaks. We generated a binarized peak accessibility count matrix for cells of each cell type, divided by genotype and gave these as input to Cicero, specifying the UMAPs generated from the LSI reductions as coordinates. For the Ngn2 dataset, Glia cells were aggregated to Progenitor clusters and Early Neurons to Glutamatergic Neurons. Additionally, for each cell type, the GADEVS and control groups of cells were down-scaled to be of the same size, as we have previously observed that the number of connections predicted by Cicero is negatively correlated with the number of input cells. Only peaks with positive accessibility correlations (Cicero score greater than zero) were included in the following analyses.

We used a custom script to find putative cis-regulatory regions for each gene, starting with co-accessibility scores. We first annotated the scATAC-seq peaks based on their overlap with promoters, exons and 5’ untranslated regions (annotation downloaded from the Ensembl database (ensembldb R package, v. 2.14.1).

We then constructed promoter-enhancer networks using the igraph R package (v. 1.2.6) for each celltype (divided by genotype), selecting only links that passed a co-accessibility threshold (0.4 for Ngn2, 0.6 for organoids).

#### Gene regulatory networks

Gene regulatory networks (GRNs) were built using CellOracle (^41^, v. 0.10.12). All peaks in the previously generated promoter-enhancer association lists were scanned for the presence of TF motifs from the gimmemotifs database (gimme.vertebrate.v5.0). Base GRNs were then generated for each combination of cell type and genotype, connecting each TF to the genes whose enhancers or promoters have the TF binding motif. Using CellOracle, we then filtered the GRNs based on the celltype-specific expression of the genes, derived from our scRNA-seq datasets. Edges in the resulting networks are drawn based on the concurrence of the following criteria: presence of the source TF’s binding motif in the gene promoter or its enhancers, co-expression of the TF and the gene, and the discernible impact of TF expression on the transcript levels of the gene, inferred by a Bagging Ridge ML model.

The GRNs were then processed using custom scripts and visualized with Cytoscape (^71^, v.3.10.0), selecting the yFiles Orthogonal layout algorithm.

## Acknowledgements

We thank all individuals affected with GADEVS and their parents for donating samples, Dr. Daniele Capocefalo for feedbacks on network analysis, we thank all European Institute of Oncology and Human Technopole facilities.

## Author contributions

M.F.P., L.R., D.A., E.T., R.S., and M.G. conducted experiments. A.V., V.F., V.A., L.M, M.R, F.D., and D.C analyzed data. M.F.P., A.V., M.G., and C.S. planned experiments. B.V., M.T.C-R., and J.F.B. provided samples. M.F.P., V.F., A.V., M.G., wrote the first draft of the manuscript. All authors contributed and approved the final manuscript. M.G, A.V, G.T. supervised and conceived the study.

## Data availability and code availability

All the data and code are available upon request.

## SUPPLEMENTARY FIGURES

**Fig. S1.**
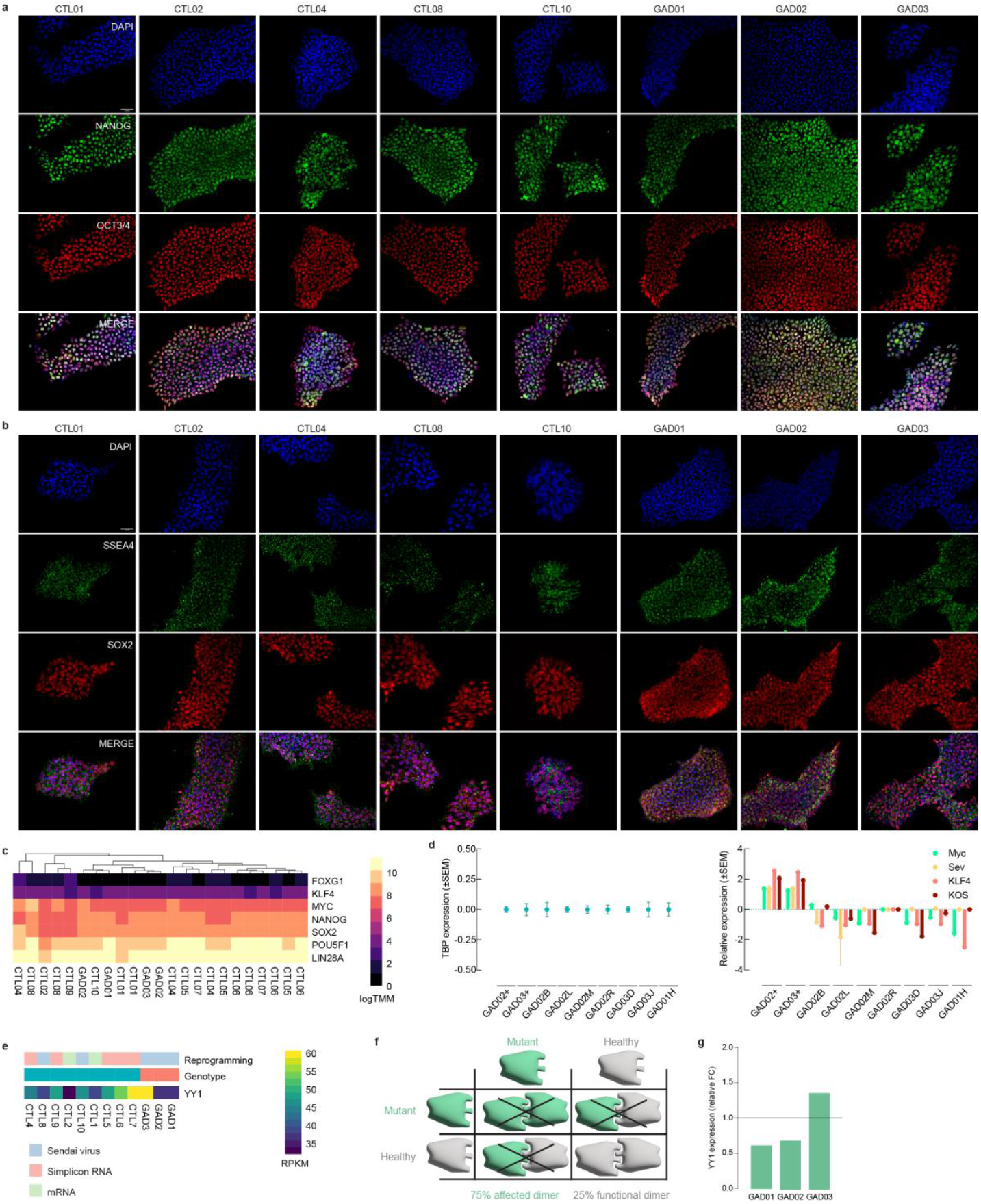
Patient-derived iPSCs display pluripotency-like features. **a**) Representative images of immunofluorescence for pluripotency markers Nanog and Oct3/4 as well as **b**) SSEA4 and Sox2 across CTL and YY1-mutated patient-derived iPSCs. Scale bar of 50 μm for all images. **c**) Newly generated cell lines express key pluripotency genes (*MYC, SOX2, NANOG, LIN28A, POU5F1*) similar to other fully established iPSC lines. Gene expression is similar across healthy and affected iPSCs. **d**) RT-qPCR targeting the viral vectors c-Myc, KLF4, KOS, SeV (and TBP for normalization) necessary for reprogramming using the Sendai virus method have been successfully removed, generating reliable vector-free clones. **e**) Heatmap showing expression values of YY1 associated with genotypes and reprogramming methods. **f**) Illustrative diagram showing YY1 dimer formation probability for GAD03 mutated iPSCs. **g**) Relative *YY1* expression in GADEVS vs. CTL samples show differential dosage in affected lines. Fold-change was calculated dividing GAD individual RPKM levels by controls average RPKM expression.

**Fig. S2.**
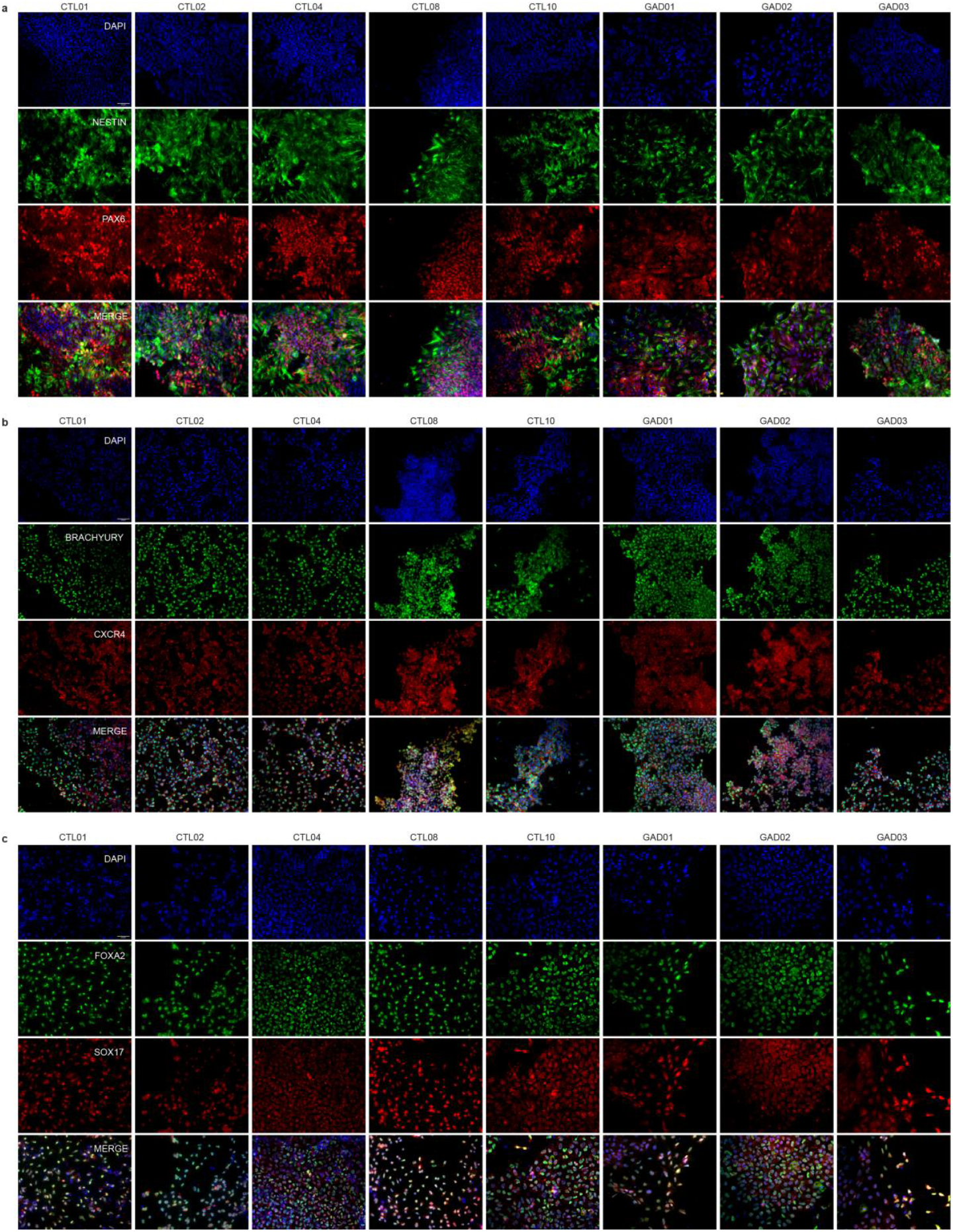
Reprogrammed iPSC lines have the potential to give rise to all three germ layer lineages. **a**) Representative images of immunocytochemistry using known markers for ectoderm (NESTIN and PAX6), **b**) mesoderm (Brachyury and CXCR4), **c**) and endoderm (FOXA2 and SOX17). Scale bar 50 μm for all images.

**Fig. S3.**
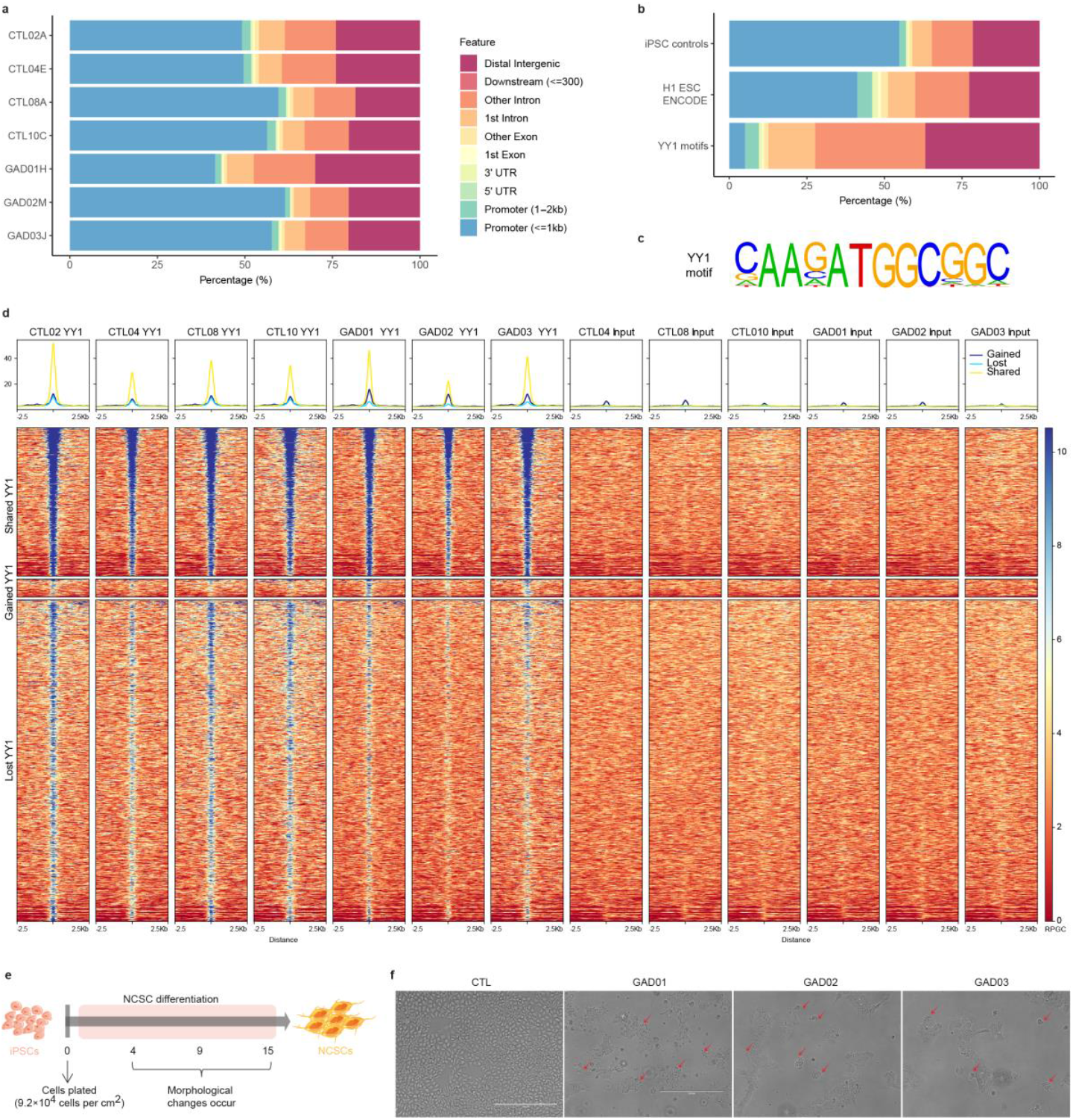
Mutated lines exhibited a global loss of binding sites in YY1 ChIPseq and fail to differentiate into NCSCs. **a**) Barplot of YY1 peaks are shown for all CTL and affected cell line. **b**) Barplot of YY1 peaks in CTLs, ENCODE ChIPseq from H1 hESC line (call sets for YY1 were downloaded from the ENCODE portal), and all YY1 motifs in the genome. **c**) Positional weighted matrix enriched of identified YY1 ChIPseq motif enrichment. **d**) YY1 ChIPseq enrichment curves are displayed for each individual CTL and GADEVS iPSC line. Upper panel represents YY1 coverage (normalized by RPKM) at gained, lost and shared peaks. Lower panel shows heatmaps (RPGC) for gained, lost and shared peaks. Each row indicates a 5 kb window centered on peak midpoints. **e**) Graphical illustration displays iPSC differentiation protocol into neural crest stem cells (NCSCs). **f**) Affected iPSCs show morphological features of cells undergoing late-stage apoptosis with red arrows pointing towards blebbing of the plasma membrane for affected lines. Images were acquired at day 14 of iPSC differentiation into NCSCs. Scale bar of 400 μm for CTL08 and 200 μm for the remaining pictures.

**Fig. S4.**
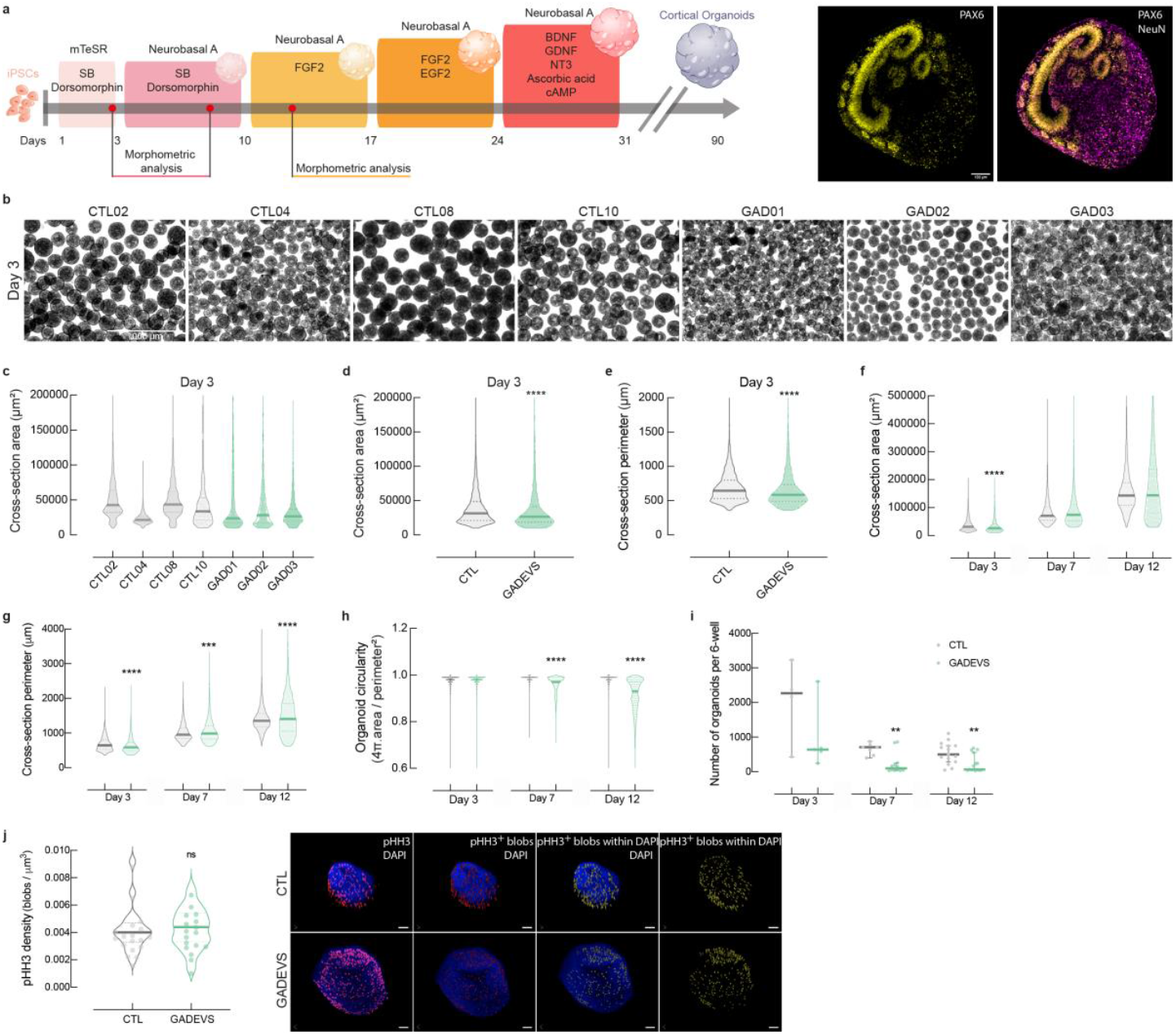
Affected lines display slight defects in circularity and generate reduced number of cortical organoids. **a**) Summarized patterning protocol used to differentiate human iPSC lines into cortical organoids. Images on the right panel are immunolabelled (PAX6 and NeuN) and cleared whole organoids at day 12 (scale bar of 100 μm). **b**) Representative figures of newly generated organoids (day 3) for CTL and GADEVS lines. Scale is 1000 μm for all pictures. **c**,**d**) Morphometric properties including cross-sectional area and **e**) perimeter are shown at day 3, as well as time-course displaying a grouped analysis for **f**) cross-sectional area, **g**) perimeter, **h**) circularity. The number of organoids per well was calculated and resulted from 2-3 batches of differentiation, with multiple wells acquired and quantified. The total number of organoids analyzed at day 3 were 12963 (*n*=8197 for CTL *vs. n*=4766 for GADEVS); day 7 were 7300 (*n*=4578 for CTL *vs. n*=2722 for GADEVS); day 12 were 11656 (*n*=8093 for CTL *vs. n*=3563 for GADEVS). **j**) pHH3 density measurements (number of blobs/ μm3) at day12 were normalized in relation to organoid volume using the DAPI channel (CTL, *n*=18; GADEVS, *n*=17). Panel on the right shows representative images of CTL and GADEVS organoids stained with DAPI and pHH3. Only pHH3 localized within DAPI is considered and counted (pHH3^+^ blobs within DAPI). Scale bar of 50 μm for all images. All data are expressed as median ± 95% confidence interval of n different organoids per experimental group. ns P>0.05, **P<0.01, ***P<0.001, ****P<0.0001, two-tailed Mann Whitney test for each individual time-point vs. CTL.

**Fig. S5.**
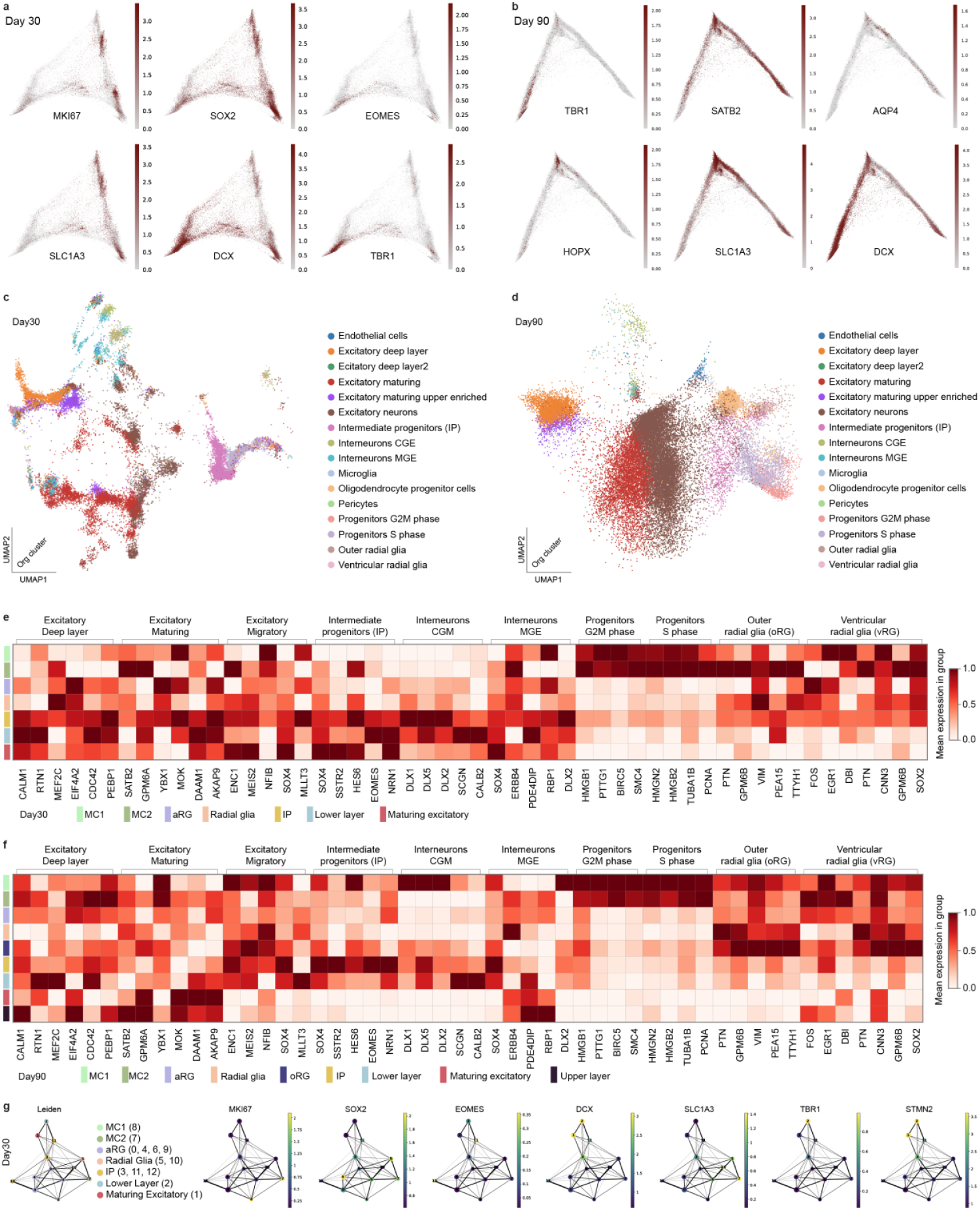
Characterization of cell-types in day 30 and day 90 cortical brain organoids. a) Diffusion maps of day 30 coloured by normalized counts for markers of cycling progenitors (MKI67), progenitors (SOX2), intermediate progenitors (EOMES), radial glia (SLC1A3), pan marker of maturing neurons (DCX), lower layer (TBR1). b) Diffusion maps of day 90 coloured by normalized counts for markers of lower layer (TBR1), upper-layer (SATB2), astrocytes (AQP4), outer radial glia/astrocytes lineage (HOPX), radial glia (SLC1A3), maturing neurons (DCX). c) UMAP derived from ingestion of day 30 organoids on a single-cell transcriptomic atlas of human neocortical development (ref: 10.1016/j.neuron.2019.06.011, corroborates the identification of populations of cycling progenitors, intermediate progenitors, maturing and lower (deep) layer neurons. d) UMAP derived from ingestion of day 90 organodis on the same single-cell transcriptomic atlas used in panel c identify a broader enrichment of excitatory neurons (TBR1^+^ in the original data) and excitatory maturing (SATB2^+^ in the original data) at later stages of cortical brain organoids differentiation; e) heatmap of mean expression per group (cell-type depicted in main figure and reported with the same colouring palette below) of genes identified by intersecting markers of organoids cell-types at day 30, with markers of cell-types identified in the atlas data used for ingestion and described for panel c-d. Expression is reported scaled from 0 (white) to 1 (red). f) heatmap of mean expression per group (cell-type depicted in main figure and reported with the same colouring palette below) of genes identified by intersecting markers of organoids cell-types at day 90, with markers of cell-types identified in the atlas data used for ingestion and described for panel c-d. Expression is reported scaled from 0 (white) to 1 (red). g) Graph of leiden-leiden relationships measured by Paga on day 30 organoids to aid cell-type identification, and respectively coloured by cell-type, and expression of key markers MKI67, SOX2, EOMES, DCX, SLC1A3, TBR1 and STMN2.

**Fig. S6.**
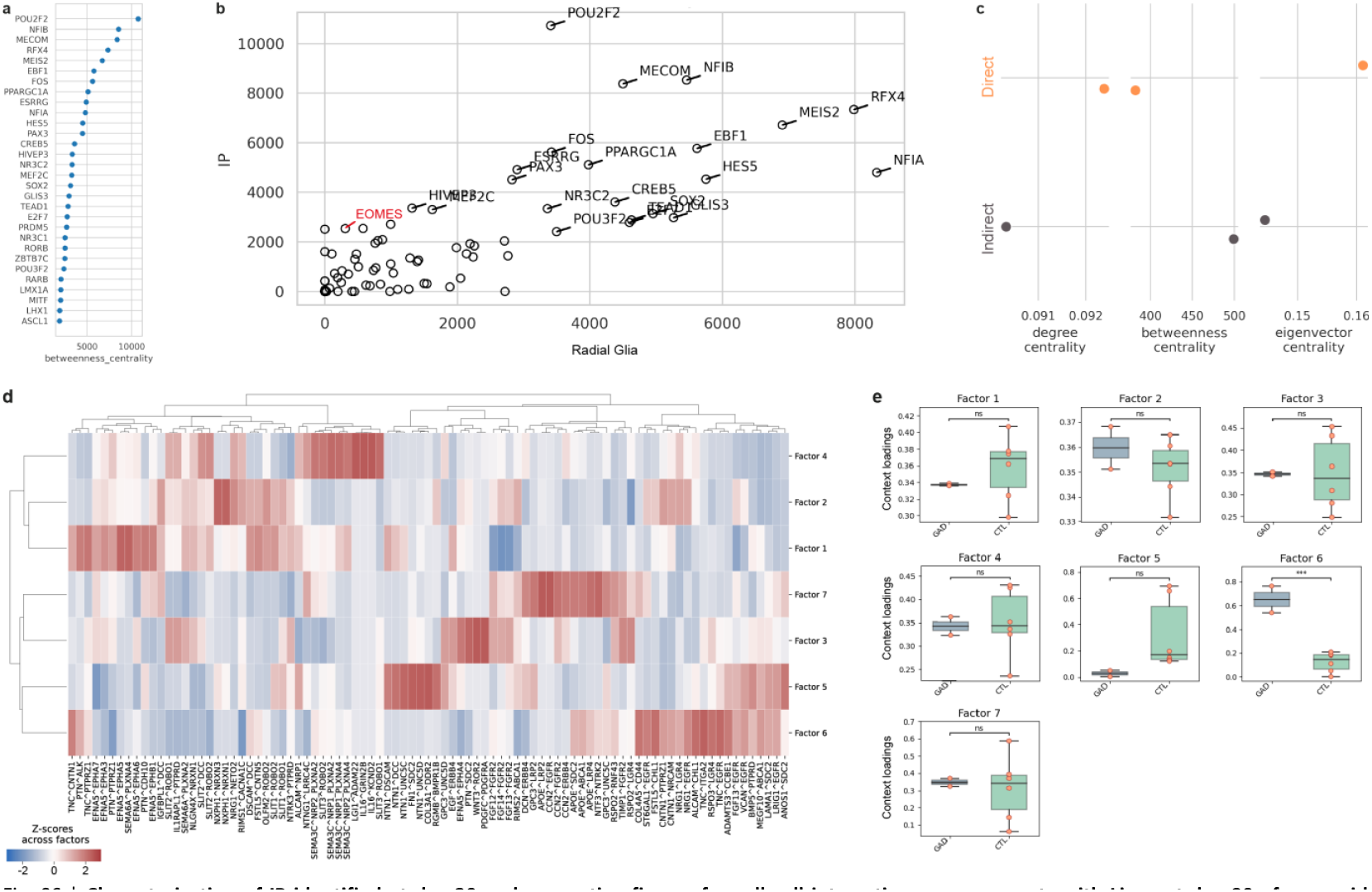
Characterization of IP identified at day 30 and supporting figures for cell-cell interaction measurements with Liana at day 90 of organoids differentiation. a) top 30 TF ranked by *betweeness centrality* represent the most active transcription factor in IP identified at day 30. b) scatter plot comparing *betweeness centrality* in IP and RG clusters identify a rather strong similarity for TFs across the two cell types. As an exclusive marker of IP, EOMES (reported in red) stands as relatively high in IP and relatively low in RG. c) Key network topology parameters calculated for EOMES in the gene-regulatory-networks (GRN) of “Direct” (IP-independent, SATB2^+^upper-layer like) and “Indirect” (IP-dependent) neuronal lineages. EOMES shows a relatively lower degree and centrality in the indirect lineages, but a higher *betweeness centrality* which points to a lower number of direct targets but a higher impact on the topology of the GRN; d) heatmap of receptor-ligand couples describing the latent factors identified by Liana in control and GADEVS organoids at day 90. Z-scores of is reported for each couple describing their contribution to each latent factor. e) boxplot of context loading for each factor.

**Fig. S7.**
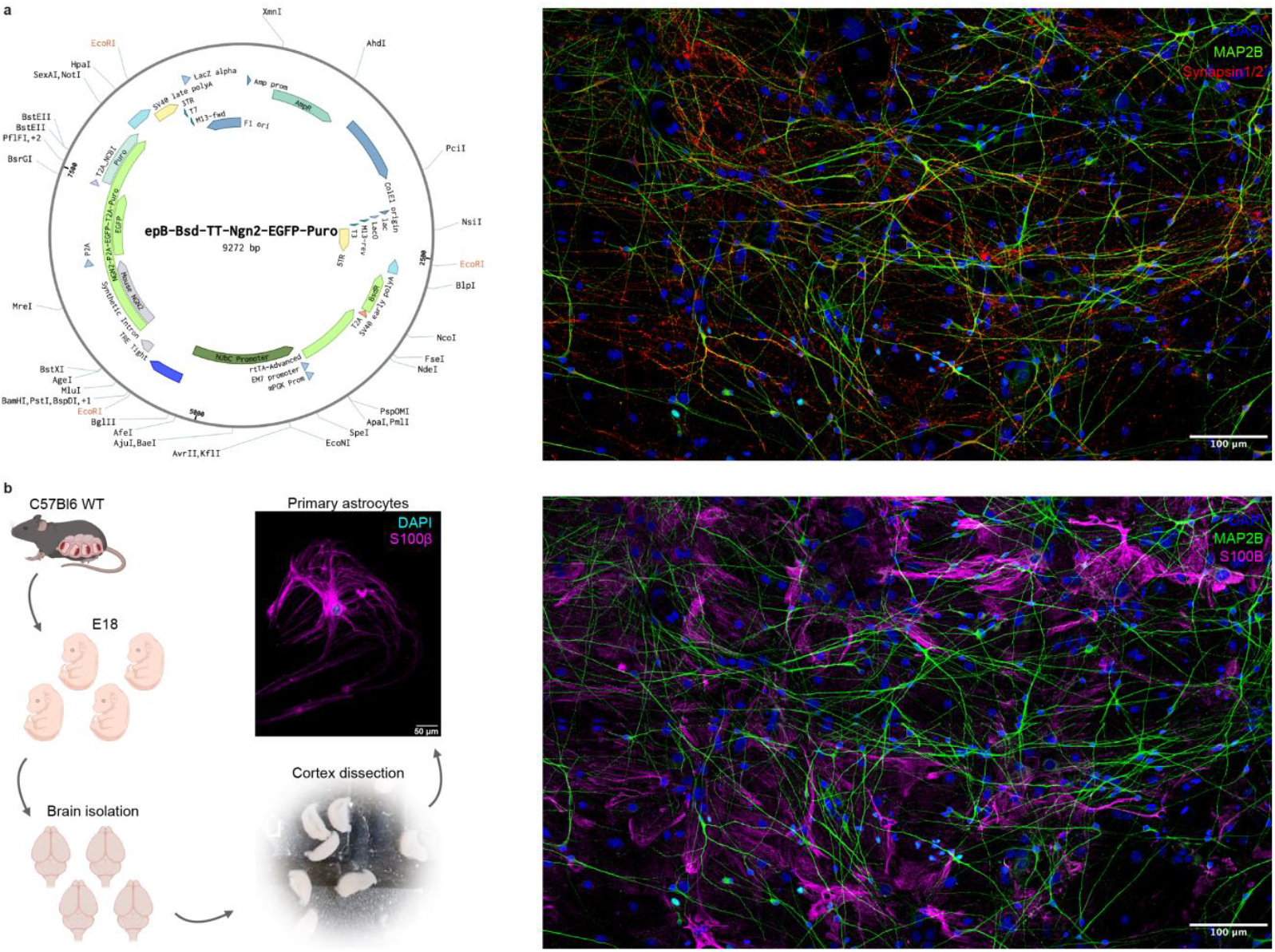
iPSC-derived glutamatergic neurons are differentiated via inducible overexpression of NGN2 and co-cultured with fresh primary mouse astrocytes. **a**) NGN2 ePB donor plasmid map used to generate glutamatergic neurons. Right panel represents iPSC-derived glutamatergic neurons immunostained with dendritic marker MAP2B and synaptic marker synapsin1/2 and acquired across multiple fields of view. **b**) Workflow overview of primary astrocyte culture generation originated from wild type mice at embryonic day 18 (E18). Right picture includes astrocytes labeled using S100β cultured with neurons. Scale bar of 100 μm.

**Fig. S8.**
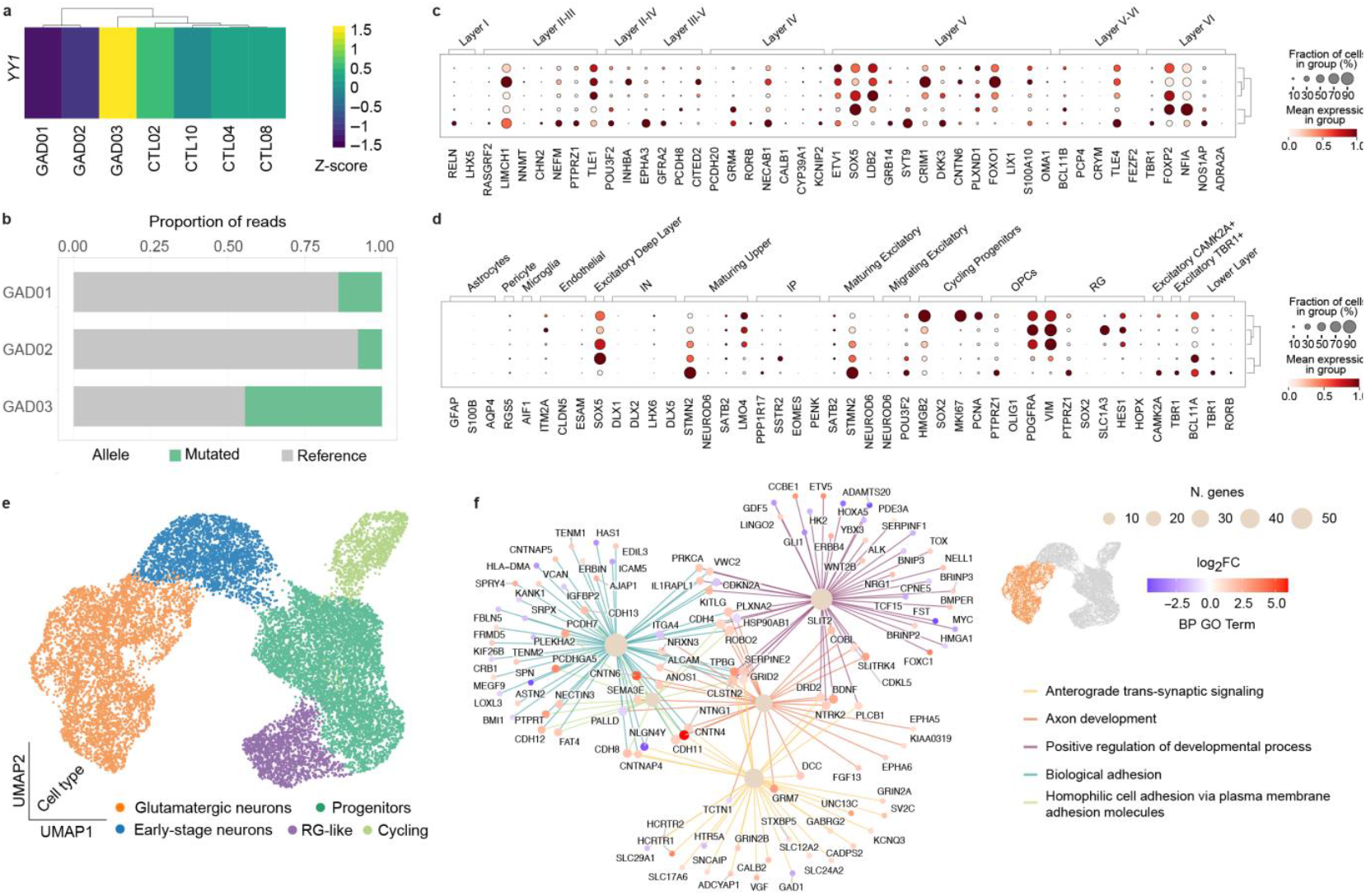
NGN2 neurons show similar levels of YY1 expression with respect to iPSC, lower layer specific marker expression and GO (biological process) enrichments similar to neurons generated in cortical brain organoids. a) Heatmap of logTMM Z-scores for the expression of YY1 in NGN2 neurons show the maintenance after differentiation of higher levels of YY1 in GAD03 and lower layer with respect to controls in GAD01 and GAD02. b) Higher coverage in NGN2 neurons allowed to quantify allele-specific gene expression in each GADEVS sample showing a 50% expression of the mutated allele only in GAD03. c) Dotplot of Layer-specific markers expression (mean expression by cell-type) in NGN2 from single-cell RNA-seq f) Dotplot of markers derived from the cortex atlas paper used for ingestion of organoids data and described for panel S5c-d: mean expression by cell-type in NGN2 single-cell RNA-seq, highlight that mature neurons bear expression of lower layer markers and CAMK2A (marker of synaptic activity/maturiity). e) UMAP of the RNA-modality of NGN2 neurons. Cell-type specific colouring is depicted on the bottom side of the panel. f) Bipartite GO-genes graph of DEGs enriching glutamatergic neurons. Nodes size for GO category is defined by the number of genes enriching them, gene nodes color are proportioned to the log2FC of differential expression measured by comparing controls and GADEVS mature neurons on pseudobulk of mature neurons. Genes referring to *positive regulation of developemntal process* include master regulators identified in fig. 6e-f MYC, ETV5 and HOXA5.

## References

1. Gabriele, M. et al. YY1 Haploinsufficiency Causes an Intellectual Disability Syndrome Featuring Transcriptional and Chromatin Dysfunction. Am. J. Hum. Genet. 100, 907–925 (2017).

2. Vissers, L. E. L. M. et al. A de novo paradigm for mental retardation. Nat. Genet. 42, 1109–1112 (2010).

3. Ferng, A., Thulin, P., Walsh, E., Weissbrod, P. A. & Friedman, J. YY1: A New Gene for Childhood Onset Dystonia with Prominent Oromandibular-Laryngeal Involvement? Mov. Disord. 37, 227–228 (2022).

4. Zorzi, G. et al. YY1-Related Dystonia: Clinical Aspects and Long-Term Response to Deep Brain Stimulation. Mov. Disord. Off. J. Mov. Disord. Soc. 36, 1461–1462 (2021).

5. Nabais Sá, M. J., Gabriele, M., Testa, G. & de Vries, B. B. Gabriele-de Vries Syndrome. in GeneReviews® (eds. Adam, M. P.et al.) (University of Washington, Seattle, Seattle (WA), 2019).

6. Woo, H., Kim, W. S. & Kim, J. S. A rare epilepsy phenotype in Gabriele-de Vries syndrome: A new case and literature review. Neurol. Asia 28, 1063–1067 (2023).

7. Brooker, S. M. & Mencacci, N. E. The expanding genetic landscape of myoclonus-dystonia syndrome: YY1 and ATP1A3 are added to the list. Parkinsonism Relat. Disord. 117, 105929 (2023).

8. Donohoe, M. E. et al. Targeted disruption of mouse Yin Yang 1 transcription factor results in peri-implantation lethality. Mol. Cell. Biol. 19, 7237–7244 (1999).

9. Verheul, T. C. J., van Hijfte, L., Perenthaler, E. & Barakat, T. S. The Why of YY1: Mechanisms of Transcriptional Regulation by Yin Yang 1. Front. Cell Dev. Biol. 8, (2020).

10. Weintraub, A. S. et al. YY1 Is a Structural Regulator of Enhancer-Promoter Loops. Cell 171, 1573-1588.e28 (2017).

11. Beagan, J. A. et al. YY1 and CTCF orchestrate a 3D chromatin looping switch during early neural lineage commitment. Genome Res. 27, 1139–1152 (2017).

12. Vitriolo, A., Gabriele, M. & Testa, G. From enhanceropathies to the epigenetic manifold underlying human cognition. Hum. Mol. Genet. 28, R226–R234 (2019).

13. Gabriele, M., Lopez Tobon, A., D’Agostino, G. & Testa, G. The chromatin basis of neurodevelopmental disorders: Rethinking dysfunction along the molecular and temporal axes. Prog. Neuropsychopharmacol. Biol. Psychiatry 84, 306–327 (2018).

14. Hsieh, T.-H. S. et al. Enhancer-promoter interactions and transcription are largely maintained upon acute loss of CTCF, cohesin, WAPL or YY1. Nat. Genet. 54, 1919–1932 (2022).

15. Carminho-Rodrigues, M. T. et al. Complex movement disorder in a patient with heterozygous YY1 mutation (Gabriele-de Vries syndrome). Am. J. Med. Genet. A. 182, 2129–2132 (2020).

16. López-Perrote, A. et al. Structure of Yin Yang 1 oligomers that cooperate with RuvBL1-RuvBL2 ATPases. J. Biol. Chem. 289, 22614–22629 (2014).

17. Gabriele, M. et al. KMT2D haploinsufficiency in Kabuki syndrome disrupts neuronal function through transcriptional and chromatin rewiring independent of H3K4-monomethylation. 2021.04.22.440945 Preprint at 10.1101/2021.04.22.440945 (2021).

18. Adamo, A. et al. 7q11.23 dosage-dependent dysregulation in human pluripotent stem cells affects transcriptional programs in disease-relevant lineages. Nat. Genet. 47, 132–141 (2015).

19. Birk, E. et al. SOBP is mutated in syndromic and nonsyndromic intellectual disability and is highly expressed in the brain limbic system. Am. J. Hum. Genet. 87, 694–700 (2010).

20. de Luis, O., Valero, M. C. & Jurado, L. A. WBSCR14, a putative transcription factor gene deleted in Williams-Beuren syndrome: complete characterisation of the human gene and the mouse ortholog. Eur. J. Hum. Genet. EJHG 8, 215–222 (2000).

21. Bauer, C. K. et al. Gain-of-Function Mutations in KCNN3 Encoding the Small-Conductance Ca2+-Activated K+ Channel SK3 Cause Zimmermann-Laband Syndrome. Am. J. Hum. Genet. 104, 1139–1157 (2019).

22. Mulley, J. C. et al. The Role of Seizure-Related SEZ6 as a Susceptibility Gene in Febrile Seizures. Neurol. Res. Int. 2011, 917565 (2011).

23. Gunnersen, J. M. et al. Sez-6 proteins affect dendritic arborization patterns and excitability of cortical pyramidal neurons. Neuron 56, 621–639 (2007).

24. Goto, S. et al. Immunocytochemical detection of calcineurin and microtubule-associated protein 2 in central neurocytoma. J. Neurooncol. 16, 19–24 (1993).

25. Hassoun, J. et al. Central neurocytoma: a synopsis of clinical and histological features. Brain Pathol. Zurich Switz. 3, 297–306 (1993).

26. Kobayashi, K. et al. Cytoskeletal protein abnormalities in patients with olivopontocerebellar atrophy--an immunocytochemical and Gallyas silver impregnation study. Neuropathol. Appl. Neurobiol. 18, 237–249 (1992).

27. Myles-Worsley, M. et al. Deletion at the SLC1A1 glutamate transporter gene co-segregates with schizophrenia and bipolar schizoaffective disorder in a 5-generation family. Am. J. Med. Genet. Part B Neuropsychiatr. Genet. Off. Publ. Int. Soc. Psychiatr. Genet. 162B, 87–95 (2013).

28. Ragge, N. K. et al. Heterozygous mutations of OTX2 cause severe ocular malformations. Am. J. Hum. Genet. 76, 1008–1022 (2005).

29. Patat, O. et al. Otocephaly-Dysgnathia Complex: Description of Four Cases and Confirmation of the Role of OTX2. Mol. Syndromol. 4, 302–305 (2013).

30. Chassaing, N. et al. OTX2 mutations contribute to the otocephaly-dysgnathia complex. J. Med. Genet. 49, 373–379 (2012).

31. Nakajima, J. et al. De novo EEF1A2 mutations in patients with characteristic facial features, intellectual disability, autistic behaviors and epilepsy. Clin. Genet. 87, 356–361 (2015).

32. Lam, W. W. K. et al. Novel de novo EEF1A2 missense mutations causing epilepsy and intellectual disability. Mol. Genet. Genomic Med. 4, 465–474 (2016).

33. Yang, J. et al. Clinical analysis of Gabriele-de Vries caused by YY1 mutations and literature review. Mol. Genet. Genomic Med. 12, e2281 (2024).

34. Varum, S. et al. Yin Yang 1 Orchestrates a Metabolic Program Required for Both Neural Crest Development and Melanoma Formation. Cell Stem Cell 24, 637-653.e9 (2019).

35. Cheroni, C. et al. Benchmarking brain organoid recapitulation of fetal corticogenesis. Transl. Psychiatry 12, 1–16 (2022).

36. Trujillo, C. A. et al. Complex Oscillatory Waves Emerging from Cortical Organoids Model Early Human Brain Network Development. Cell Stem Cell 25, 558-569.e7 (2019).

37. Chen, S., Lake, B. B. & Zhang, K. High-throughput sequencing of the transcriptome and chromatin accessibility in the same cell. Nat. Biotechnol. 37, 1452–1457 (2019).

38. Zhu, C. et al. An ultra high-throughput method for single-cell joint analysis of open chromatin and transcriptome. Nat. Struct. Mol. Biol. 26, 1063–1070 (2019).

39. Cao, J. et al. Joint profiling of chromatin accessibility and gene expression in thousands of single cells. Science 361, 1380–1385 (2018).

40. Polioudakis, D. et al. A Single-Cell Transcriptomic Atlas of Human Neocortical Development during Midgestation. Neuron 103, 785-801.e8 (2019).

41. Kamimoto, K. et al. Dissecting cell identity via network inference and in silico gene perturbation. Nature 614, 742–751 (2023).

42. Moriano, J., Leonardi, O., Vitriolo, A., Testa, G. & Boeckx, C. A multi-layered integrative analysis reveals a cholesterol metabolic program in outer radial glia with implications for human brain evolution. 2023.06.23.546307 Preprint at 10.1101/2023.06.23.546307 (2023).

43. Cheon, S. Y. Impaired Cholesterol Metabolism, Neurons, and Neuropsychiatric Disorders. Exp. Neurobiol. 32, 57–67 (2023).

44. Dimitrov, D. et al. Comparison of methods and resources for cell-cell communication inference from single-cell RNA-Seq data. Nat. Commun. 13, 3224 (2022).

45. Lin, H.-C. et al. NGN2 induces diverse neuron types from human pluripotency. Stem Cell Rep. 16, 2118–2127 (2021).

46. Moulson, A. J., Squair, J. W., Franklin, R. J. M., Tetzlaff, W. & Assinck, P. Diversity of Reactive Astrogliosis in CNS Pathology: Heterogeneity or Plasticity? Front. Cell. Neurosci. 15, (2021).

47. Escartin, C. et al. Reactive astrocyte nomenclature, definitions, and future directions. Nat. Neurosci. 24, 312–325 (2021).

48. Matusova, Z., Hol, E. M., Pekny, M., Kubista, M. & Valihrach, L. Reactive astrogliosis in the era of single-cell transcriptomics. Front. Cell. Neurosci. 17, 1173200 (2023).

49. Abd-El-Basset, E. M., Rao, M. S., Alshawaf, S. M., Ashkanani, H. K. & Kabli, A. H. Tumor necrosis factor (TNF) induces astrogliosis, microgliosis and promotes survival of cortical neurons. AIMS Neurosci. 8, 558–584 (2021).

50. Girton, J. R. & Jeon, S. H. Novel embryonic and adult homeotic phenotypes are produced by pleiohomeotic mutations in Drosophila. Dev. Biol. 161, 393–407 (1994).

51. Kwon, H.-J. & Chung, H.-M. Yin Yang 1, a vertebrate polycomb group gene, regulates antero-posterior neural patterning. Biochem. Biophys. Res. Commun. 306, 1008–1013 (2003).

52. Morgan, M. J., Woltering, J. M., In der Rieden, P. M. J., Du ton, A. J. & Thiery, J. P. YY1 regulates the neural crest-associated slug gene in Xenopus laevis. J. Biol. Chem. 279, 46826–46834 (2004).

53. Zurkirchen, L. et al. Yin Yang 1 sustains biosynthetic demands during brain development in a stage-specific manner. Nat. Commun. 10, 2192 (2019).

54. Dong, X. & Kwan, K. M. Yin Yang 1 is critical for mid-hindbrain neuroepithelium development and involved in cerebellar agenesis. Mol. Brain 13, 104 (2020).

55. Khamirani, H. J. et al. Clinical features of patients with Yin Yang 1 deficiency causing Gabriele-de Vries syndrome: A new case and review of the literature. Ann. Hum. Genet. 86, 52–62 (2022).

56. Schildge, S., Bohrer, C., Beck, K. & Schachtrup, C. Isolation and culture of mouse cortical astrocytes. J. Vis. Exp. JoVE 50079 (2013) doi:10.3791/50079.

57. Pereira, M. F. et al. l-α-aminoadipate causes astrocyte pathology with negative impact on mouse hippocampal synaptic plasticity and memory. FASEB J. Off. Publ. Fed. Am. Soc. Exp. Biol. 35, e21726 (2021).

58. Singh, R. et al. Autism Spectrum Disorders: A Recent Update on Targeting Inflammatory Pathways with Natural Anti-Inflammatory Agents. Biomedicines 11, 115 (2023).

59. Członkowska, A. & Kurkowska-Jastrzębska, I. Inflammation and gliosis in neurological diseases – clinical implications. J. Neuroimmunol. 231, 78–85 (2011).

60. Çarçak, N., Onat, F. & Sitnikova, E. Astrocytes as a target for therapeutic strategies in epilepsy: current insights. Front. Mol. Neurosci. 16, (2023).

61. Coulter, D. A. & Steinhäuser, C. Role of astrocytes in epilepsy. Cold Spring Harb. Perspect. Med. 5, a022434 (2015).

62. Lawrence, J. M., Schardien, K., Wigdahl, B. & Nonnemacher, M. R. Roles of neuropathology-associated reactive astrocytes: a systematic review. Acta Neuropathol. Commun. 11, 42 (2023).

63. Menendez, L. et al. Directed differentiation of human pluripotent cells to neural crest stem cells. Nat. Protoc. 8, 203–212 (2013).

64. Zhang, Y. et al. Rapid Single-Step Induction of Functional Neurons from Human Pluripotent Stem Cells. Neuron 78, 785–798 (2013).

65. Patro, R., Duggal, G., Love, M. I., Irizarry, R. A. & Kingsford, C. Salmon provides fast and bias-aware quantification of transcript expression. Nat. Methods 14, 417–419 (2017).

66. Zanella, M. et al. Dosage analysis of the 7q11.23 Williams region identifies BAZ1B as a major human gene patterning the modern human face and underlying self-domestication. Sci. Adv. 5, (2019).

67. Alexa, A., Rahnenführer, J. & Lengauer, T. Improved scoring of functional groups from gene expression data by decorrelating GO graph structure. Bioinformatics 22, 1600–1607 (2006).

68. Wu, T. et al. clusterProfiler 4.0: A universal enrichment tool for interpreting omics data. Innov. Camb. Mass 2, 100141 (2021).

69. Tan, G. & Lenhard, B. TFBSTools: an R/bioconductor package for transcription factor binding site analysis. Bioinforma. Oxf. Engl. 32, 1555–1556 (2016).

70. Garcia-Alonso, L., Holland, C. H., Ibrahim, M. M., Turei, D. & Saez-Rodriguez, J. Benchmark and integration of resources for the estimation of human transcription factor activities. Genome Res. 29, 1363–1375 (2019).

71. Shannon, P. et al. Cytoscape: a software environment for integrated models of biomolecular interaction networks. Genome Res. 13, 2498–2504 (2003).

72. Ramírez, F., Dündar, F., Diehl, S., Grüning, B. A. & Manke, T. deepTools: a flexible platform for exploring deep-sequencing data. Nucleic Acids Res. 42, W187–191 (2014).

73. Caporale, N. et al. Multiplexing cortical brain organoids for the longitudinal dissection of developmental traits at single cell resolution. 2023.08.21.553507 Preprint at 10.1101/2023.08.21.553507 (2023).

74. Wolf, F. A., Angerer, P. & Theis, F. J. SCANPY: large-scale single-cell gene expression data analysis. Genome Biol. 19, 15 (2018).

75. Hunter, J. D. Matplotlib: A 2D Graphics Environment. Comput. Sci. Eng. 9, 90–95 (2007).

76. Waskom, M. L. seaborn: statistical data visualization. J. Open Source Softw. 6, 3021 (2021).

77. McKinney, W. Data Structures for Statistical Computing in Python. Proc. 9th Python Sci. Conf. 56–61 (2010) doi:10.25080/Majora-92bf1922-00a.

78. Harris, C. R. et al. Array programming with NumPy. Nature 585, 357–362 (2020).

79. Robinson, M. D., McCarthy, D. J. & Smyth, G. K. edgeR: a Bioconductor package for differential expression analysis of digital gene expression data. Bioinformatics 26, 139–140 (2010).

80. Gene Ontology Consortium. The Gene Ontology resource: enriching a GOld mine. Nucleic Acids Res. 49, D325–D334 (2021).

81. Granja, J. M. et al. ArchR is a scalable software package for integrative single-cell chromatin accessibility analysis. Nat. Genet. 53, 403–411 (2021).

82. Wickham, H. Ggplot2. (Springer International Publishing, Cham, 2016). doi:10.1007/978-3-319-24277-4.

83. Zhang, Y. et al. Model-based Analysis of ChIP-Seq (MACS). Genome Biol. 9, R137 (2008).

84. Castro-Mondragon, J. A. et al. JASPAR 2022: the 9th release of the open-access database of transcription factor binding profiles. Nucleic Acids Res. 50, D165– D173 (2022).

85. Schep, A. N., Wu, B., Buenrostro, J. D. & Greenleaf, W. J. chromVAR: inferring transcription-factor-associated accessibility from single-cell epigenomic data. Nat. Methods 14, 975–978 (2017).

86. Pliner, H. A. et al. Cicero Predicts cis-Regulatory DNA Interactions from Single-Cell Chromatin Accessibility Data. Mol. Cell 71, 858-871.e8 (2018).

